# Timelapse viability assay to detect division and death of primary multiple myeloma cells in response to drug treatments with single cell resolution

**DOI:** 10.1101/2021.11.12.467843

**Authors:** Christina Mark, Natalie S. Callander, Kenny Chng, Shigeki Miyamoto, Jay Warrick

**Author notes:** Salus Discovery, LLC, 110 E. Main St. Suite 815, Madison, WI 53703.

## Abstract

Heterogeneity among cancer cells and in the tumor microenvironment (TME) is thought to be a significant contributor to the heterogeneity of clinical therapy response observed between patients and can evolve over time. A primary example of this is multiple myeloma (MM), a generally incurable cancer where such heterogeneity contributes to the persistent evolution of drug resistance. However, there is a paucity of functional assays for studying this heterogeneity in patient samples or for assessing the influence of the patient TME on therapy response. Indeed, the population-averaged data provided by traditional drug response assays and the large number of cells required for screening remain significant hurdles to advancement. To address these hurdles, we developed a suite of accessible technologies for quantifying functional drug response to a panel of therapies in *ex vivo* three-dimensional culture using small quantities of a patient’s own cancer and TME components. This suite includes tools for label-free single-cell identification and quantification of both cell division and death events with a standard brightfield microscope, an open-source software package for objective image analysis and feasible data management of multi-day timelapse experiments, and a new approach to fluorescent detection of cell death that is compatible with long-term imaging of primary cells. These new tools and capabilities are used to enable sensitive, objective, functional characterization of primary MM cell therapy response in the presence of TME components, laying the foundation for future studies and efforts to enable predictive assessment drug efficacy for individual patients.

**Insight Box:** The new tools and capabilities described here allow new insights into functional primary cell therapy response by (i) enabling more drugs to be tested on precious and limited numbers of patient cells *ex vivo* (ii) providing the ability to discriminate both cell division and death events over multiple days with single-cell resolution, and (iii) by incorporating the influences of a patient’s own cancer cells and TME components on drug response.

## Introduction

Heterogeneity in cancer cells within individual cancer patients has been well documented through single-cell “omics” analyses, such as single cell RNA sequencing and single cell proteomics, using primary patient samples.^1–4^As patients undergo therapies, such single cell omics approaches are uncovering the evolution of cancer cell clones with different fitness.^5,6^ While single cell omics help define the complexity and heterogeneity of primary human cancer cells, they do not directly measure functional behaviors of individual cells. Moreover, cancer cell behaviors can be significantly modified by the presence of other cell types and soluble and insoluble factors in the tumor microenvironment (TME), which further complicates the ability to extract tumor cell behavioral information, such as drug response or resistance, from single cell omics approaches.^7,8^ Thus, an ability to define biological behaviors of primary cancer cells at the single cell level could complement single cell omics approaches to understand the evolving human cancer cell heterogeneity in individual patients.

Functional analysis of primary cancer cells poses particularly difficult challenges due to the frequently limited number of cancer cells that are available for analysis. In addition, classical assays that are used to analyze cancer cell behaviors and drug responses, such as MTT, BrdU incorporation, clonogenic survival, apoptosis and other viability assays, generally require termination of cells to obtain data points. Thus, acquisition of statistically amenable data requires multiple parallel assays with therapeutic drugs at an arbitrary time point. If one wishes to obtain information regarding kinetics of drug responses, multiple time points are required, a situation that is often infeasible with limited primary patient cells. Moreover, these assays provide population cell averages and do not provide single cell resolution in their readouts. Thus, the development of cell assays that enable the functional analysis of individual primary cancer cells at multiple time points without requiring their termination would help advance the understanding of human cancer cell heterogeneity through *functional* endpoints. It is further desirable that such assays have the capacity to incorporate a patient’s unique TME components to define their influence on cancer cell behaviors. In the present study, we aimed to develop a time-lapse cell imaging assay that is capable of quantifying key cellular behaviors, such as division and death, with single cell resolution and robust statistical analysis. We employed a multiple myeloma (MM) cell line as well as patient samples to develop our assay.

MM is characterized by the infiltration of malignant plasma cells often at multiple sites in the bone marrow.^9–11^ There are an array of therapeutic options for treatment of MM, such as proteasome inhibitors and others; however, MM is generally considered incurable and remains the second most common blood cancer in the United States.^9–13^ MM is a heterogeneous disease in its genetic makeup, clinical manifestations, and drug responses. Moreover, drug response is influenced by multiple components of the TME, including cells (e.g., bone marrow stromal cells (BMSCs)), extracellular matrix (ECM), and soluble factors.^14–16^ To analyze responses of primary MM cells *ex vivo*, multiple assays have been developed with or without TME components.^17–20^ These assays utilize an exclusion dye to identify viable cells, colorimetric assays to measure cell metabolic activity, clonogenic assays, or 3-D culture system with TME cells to mimic the MM niche.^21^ The notable hurdles with the functional analysis of MM primary cancer cells *ex vivo* include the tendency of primary MM cells being often prone to stress and apoptosis once isolated from bone marrow aspirate, the variable quantity of MM malignant cells that can be extracted from each patient bone marrow aspirate, and the heterogeneous nature of each patient disease stage, current therapy regimen, and tumor ecosystem. To address these challenges, we previously developed a microchannel-based cis-co-culture (µC3) system that enables cytotoxicity assay for MM cells in the presence of the patient’s unique tumor-associated TME cells.^22^ While we reported a correlation of cell viability in µC3 assay platform to bortezomib, a clinically used proteasome inhibitor, with clinical responses of individual patients to bortezomib-containing therapy, the assay also posed limitations. The culture wells were limited to interaction via soluble factors and toxicity was assessed with dose curves at a single time point, which quickly depletes valuable patient cell samples. Moreover, the data collection required termination of cultures and the resulting data were not at the single cell level but rather cell population averages. Finally, the *ex vivo* health of primary cells without any drug treatment was variable and not robust, thus requiring data normalization to drug-naive control groups.

To address these shortcomings, we developed a time-lapse drug sensitivity assay methodology to more cohesively capture the behaviors of individual drug-treated primary MM cells *ex vivo*. The method utilizes an automated, inverted fluorescence microscope equipped with on-stage incubator (Figure 1). The MM culture contains 3D collagen to mimic the myeloma extracellular matrix to help provide cues typically seen in the TME such as organization and stiffness of the supporting extracellular matrix, as described by Khin et al.^23^ To study the impacts of additional TME components, we have incorporated patient-matched bone marrow plasma and non-MM bone marrow cell fractions in this assay. Lastly, we developed a sensitive method for detecting cell division and cell death events in real-time using a live-cell compatible DNA dye and quantitative phase imaging (QPI). QPI enables quantification of time-dependent changes in cellular shape and density/permeability without staining, harvesting, or labeling the cells.^24^

**Figure 1.**
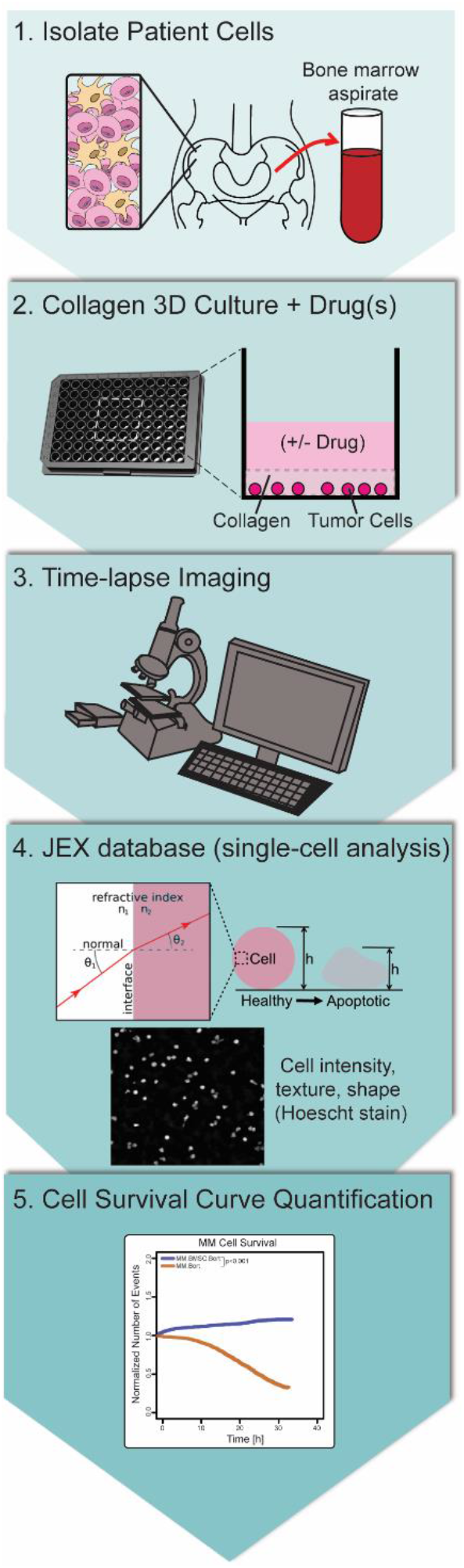
Live cell imaging time-lapse workflow. Primary cells are isolated from multiple myeloma bone marrow aspirate in 1 to be cultured in a layer of collagen and seeded in a 96-well plate in the presence of drugs in 2. Images are taken via microscopy for 72 hours in 3 and then loaded into a single-cell analysis software in 4. The software produces an R plug-in table from the dead cells that are counted using specific metrics, and the table becomes an input to analysis survival curves via R Studio software in 5.

We also developed the open-source JEX software package to provide objective image analysis and feasible data management of large timelapse experiments using a desktop computer.^25^ These advancements set the stage for more advanced functional assessments with single cell resolution (e.g., incorporating image-based machine learning) in the future to complement currently employed single cell omics studies.

## Results/discussion

We developed a timelapse imaging assay for functional characterization of limited amounts of primary patient cells. Key technical components of the assay are described first, followed by characterization of performance with a MM cell line and, finally, application of the method to characterize multi-drug responses in primary cells.

### Key Technical Components of the Assay

Three key technical components of the system that were developed include: (i) a new open-source, Java-based implementation of transport-of-intensity equation (TIE) for quantitative phase imaging (QPI) with a novel and computationally efficient method of noise reduction for robust label-free single-cell identification and detection of apoptosis events; (ii) a method for using Hoechst 33258 as a biochemical marker of cell death to provide enhanced detection of apoptosis events while minimizing potential perturbation of cells during timelapse imaging; and (iii) the JEX software package to provide objective image analysis and feasible data management of large timelapse experiments using a desktop computer.

#### Advanced implementation of TIE-QPI

QPI is the practice of quantifying phase shifts in light (Δφ) as it passes through a sample at every pixel location within an image. These phase shifts are valueable in the context of live-cell imaging because they can be measured without using labels, and they can be directly related to physical characteristics of the cells using the simple equation Δφ = d*Δn, where *d* is the cell thickness and *n* is the relative index of refraction compared to the surrounding medium.

In the case of cells, index of refraction is dictated by the density (i.e., protein / water content). Although there are many ways to perform QPI, the transport-of-intensity equation (TIE) is the only approach that can be used with a standard brightfield microscope.^24^ For these reasons, we chose to implement TIE-QPI to facilitate label-free quantification of primary cell drug response.

One key technical advancement provided by work is the open-source, Java-based implementation of QPI. Our data analysis pipeline relies heavily on the use of Java libraries, yet we could not find a Java-based implementation of the TIE-QPI. Therefore, TIE was integrated into our platform by translating Matlab code provided by Zou et al.^26^ into Java for use with well-established ImageJ Java libraries and is now available to the research community.

We also developed a novel approach to address the primary weakness of TIE-QPI that has limited its use more broadly. The primary weakness of TIE-QPI for biological applications and real-world image data is the inherent presence of image artifacts that arise from limited knowledge of boundary conditions and from low frequency noise introduced during Fourier transform-based integration steps.^27^ Given the severity of these artifacts, use of TIE-QPI to study cells has been limited to a small number of studies with small numbers of images.^24^ The current standard approach to suppressing these artifacts is to use a high-pass frequency filter to remove low frequency background variations and retain finer foreground details like cells (see ESI). This frequency filter is simple, ubiquitous, and integrates seamlessly with the Fourier-based calculation methods used for TIE-QPI. However, we found that variations and processing artifacts to be prohibitive for accurate long-term timelapse analysis of single cells (e.g., dark and light spots indicated with arrows in Figure 3F), so alternative approaches to filtering these artifacts were needed. Traditional methods like rolling-ball background subtraction were not able to deal with dark image artifacts left by TIE-QPI and low frequency filtering. A rolling ball filter determines the floor of the image background rather than the mean of the background. Therefore after subtraction, a highly variable amount of background signal is left behind that prevent use of uniform analysis parameters downstream. Median-based filtering finds the mean of the noise but is too computationally time-consuming for the large datasets, even when using computationally optimized implementations. Standard mean-based background subtraction is fast, but was far too ineffective. Therefore, we implemented a new weighted-mean background subtraction method that leverages differences in the pixel-wise variance seen in background pixels compared to objects of interest. This method is able to robustly deal with the TIE-QPI image artifacts, performed well in clusters of objects that typically pose a challenge to kernel based techniques, and is both fast and robust enough to quantitatively treat the large timelapse datasets. A comparison of the new method with rolling-ball and median background subtraction is shown in Figure 2 using a fluorescent image where the absorbance of light by red blood cells creates image features that are well below normal background, thereby creating challenges for some background subtraction methods. Figure 3 shows the superior performance of this method when used to enhance noise/background reduction in TIE-QPI compared to the standard FFT-based filter. The stark difference in the uniformity of the background and ability to handle TIE-QPI image artifacts (arrows) is clear when comparing the insets of Figure 3E and 3F. A further benefit of the approach is that it can also be used to simultaneously perform illumination correction in fluorescence images (via division of the signal by background levels) as well as generate threshold masks (via thresholding of the weighting mask), as described in the ESI. The weighted-mean background correction algorithm was instrumental in enabling robust background subtraction of results across all datasets with no changes in parameters for consistent results and objective automated analysis. See the ESI for algorithm details.

**Figure 2.**
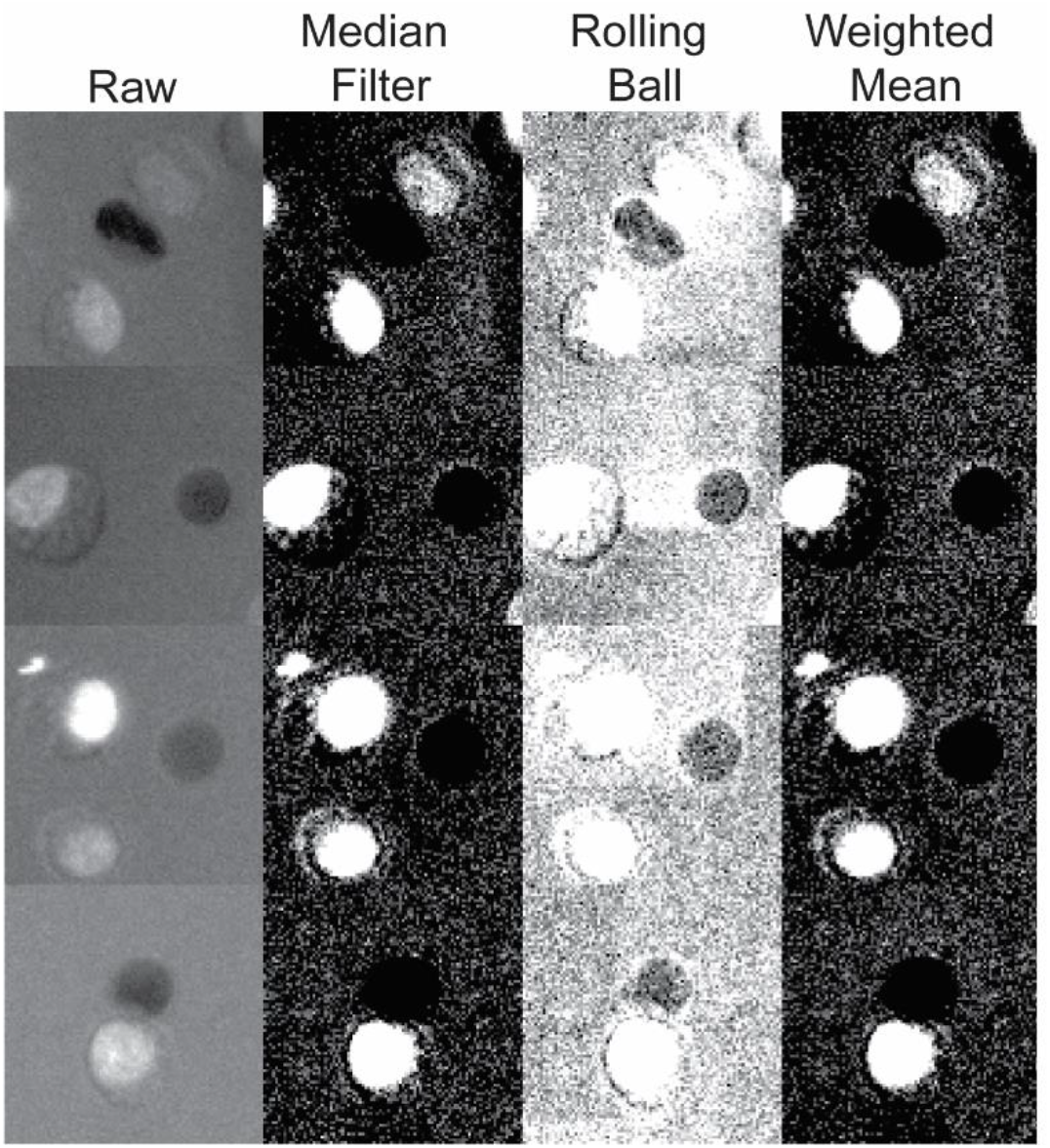
Algorithm for enhanced background subtraction. Images in the last 3 columns are contrasted identically for accurate comparison. The width of each image tile is 30 µm. Contrast is enhanced to illustrate the artifacts that affect subsequent image segmentation and cell identification.

**Figure 3.**
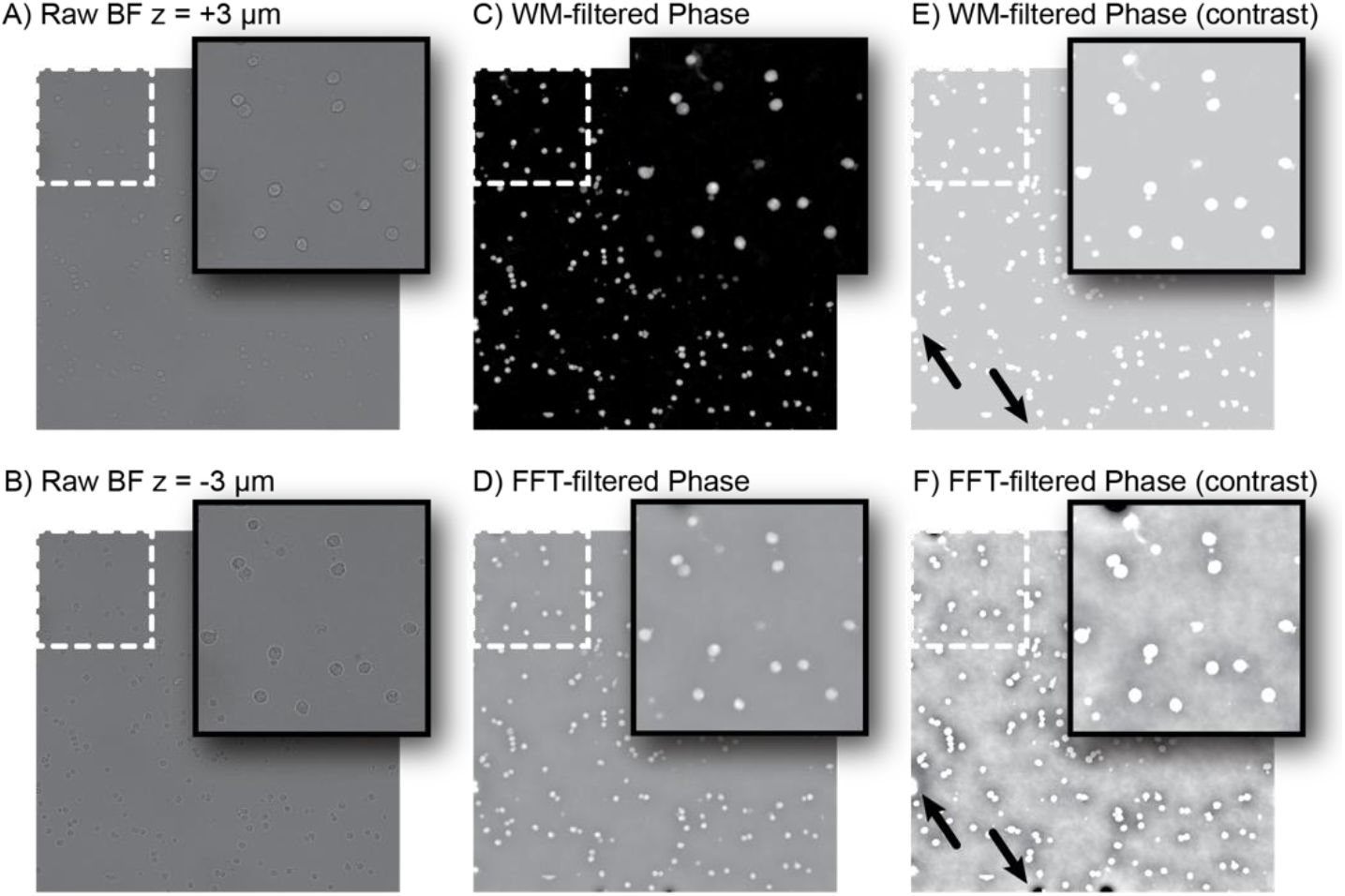
Quantitative phase imaging (QPI). (A) Raw brightfield image of patient MM cells at the z = +3 µm plane. (B) Raw brightfield image of patient MM cells at the z = - 3 µm plane. Notice the different intensities within cells compared to the z = +3 µm plane, which is measured by the ∂I/∂z in TIE equation (ESI). (C) Result of the TIE-QPI computation (Δφ) after application of the new noise filtering approach that leverages noise filtering approach that leverages weighted-mean filtering (WM-filtering). (D) Result of the TIE-QPI computation (Δφ) using just the traditional FFT-filtering method (see ESI Eq 3 for details). (E) A contrast enhanced version of the image in subfigure D. (F) The image of subfigure E contrast enhanced identically to the image in subfigure E for direct comparison of background at identical gains and offsets.

#### Live cell nuclear staining to detect cell permeability upon death

The second major advancement of this work is a new method for using Hoechst 33258 live-cell nuclear stain (vs. the more common Hoechst 33342) as an indicator of apoptosis in timelapse microscopy. The goal of this imaging and labeling method was to measure cell permeability while perturbing the cells as little as possible during imaging by (i) limiting the concentration of dye used in the culture media during timelapse and (ii) avoiding DNA damage directly from UV light and/or indirectly from reactive oxygen species (ROS) that can be produced by fluorescent dyes during illumination. In terms of dye concentration, Hoechst 33258 has been previously shown to not induce apoptosis at relatively high doses (∼ 26.7 µM for 24 hours) in multiple cell lines^28,29^. Here we use Hoechst 33258 at 0.64 µM (41-fold lower concentration). Previous papers have also shown that lengthening the wavelength of UV illumination from 365 nm to 405 nm can reduce degree of DNA damage from extended exposures (10-300 s) by 10-25-fold.^30^ Literature recommends using less than 2.6 J/cm^2^ total dose of 365 nm UV light during timelapse microscopy.^31^ Here we use a total dose of ∼0.8 J/cm^2^ over 48 hours of less harmful 395 nm light. Further, Hoechst 33258 exhibits 10 times less cell permeability than its more common counterpart Hoechst 33342. Therefore, at our low concentrations, image pixels in the nuclei of live cells are only ∼2 standard deviations above the background noise, indicating a low concentration of dye within live cells. By minimizing dye concentrations within the live cells, production of ROS that could harm cells during imaging is also minimized. Indeed, we found that the Hoechst labeling approach, combined with the use of the collagen overlay, eliminated loss of cell viability in vehicle-treated (untreated control) patient samples compared to previous work that experienced ∼40-70% cell loss in the first day of culture^22^ (see patient sample analysis below). Furthermore, when the cell membrane is compromised during apoptosis, fluorescent dye readily enters the cell, causing a marked increase in fluorescence (Figure 4), providing the ability to quantify single-cell death.

**Figure 4.**
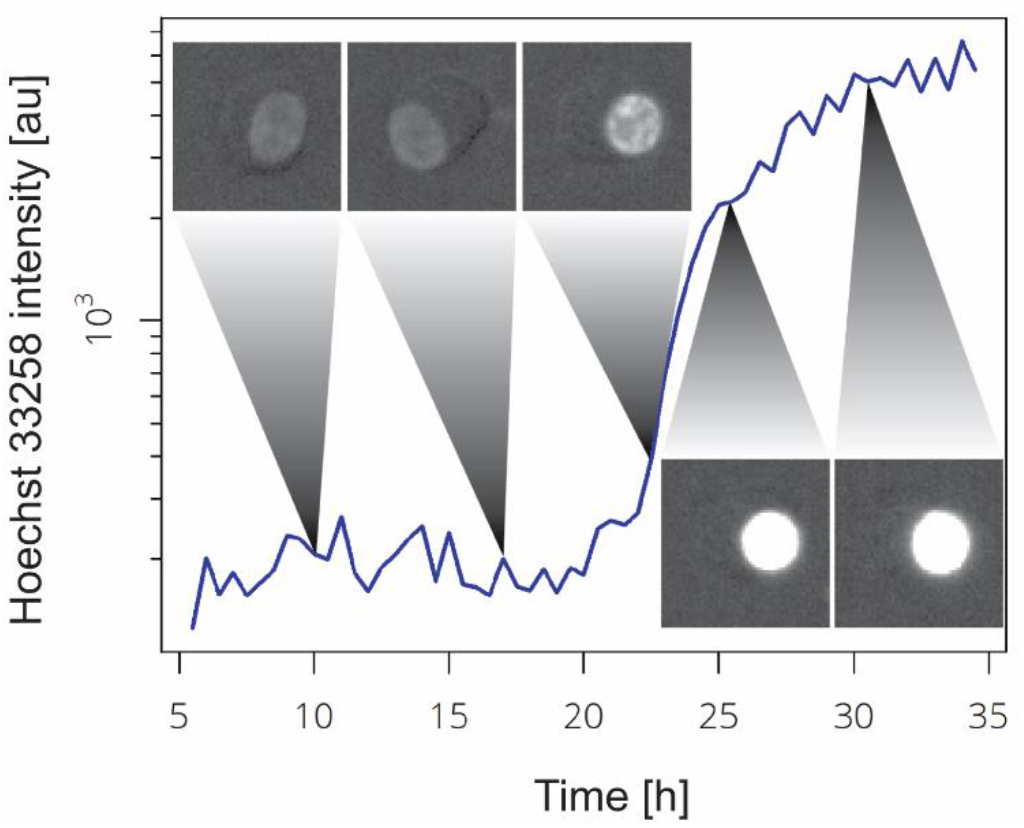
Hoechst 33258 as a viability dye for live-patient-cell timelapse. Plot of Hoechst intensity vs. time for an example cell undergoing apoptosis. Thumbnail images at equal contrast.

However, with very low live-cell nuclear staining intensities, use of nuclear fluorescence for cell counting and cell tracking was not sufficiently reliable. In contrast, the TIE-QPI phase images provided exquisite label free signals that extended to the boundary of the cell (i.e., covering both the nucleus and cytoplasm). Thus, the phase images were used to perform cell identification, masking, segmentation and tracking for quantification of both phase image intensity and Hoechst fluorescence.

Lastly, Hoechst staining intensity (HSI) and phase image intensity (Δφ) were leveraged together to provide a more sensitive and robust measure of cell death. While both metrics are aimed at detecting the same event, they are independent measures with independent noise characteristics. Therefore, both metrics were combined to provide a more robust and sensitive metric of cell viability. The final <u>v</u>iability <u>m</u>etric (VM) is calculated as VM = HSI/Δφ. A ratio was used because, when a cell dies, the Hoechst staining intensity (HSI) increases while the phase image intensity (Δφ) decreases. Therefore, the VM increases significantly upon cell death. See ESI for algorithm details.

#### JEX software for batch image analysis and data management

Finally, a major hurdle of data management and analysis was addressed using JEX software. JEX is an open-source, platform-independent Java program for batch analysis of image data.^25^ The software was critical to organizing and maintaining the large amounts of timelapse image data. When performing timelapse imaging over multiple wells (16), at multiple locations within each well (9), at many timepoints (every 30 minutes for multiple days), in multiple channels (2) and z positions (2); raw data from a single experiment is on the order of 300 Gb. Thus, standard computer RAM, ROM, processers, and file indexing become limiting factors. For example, the simple approach of putting all the images into a single folder makes subsequent data analysis intractable. When the number of files reaches ∼10,000 files, each time a file is opened by name, a hard-disk-based file system must search for 1-3 seconds to find the file before opening. Aggregated over 10,000 files results in ∼5 hours of delays for a single look-up operation in addition to the time to open and process the file. Furthermore, such delays are super-linear with file number, rapidly creating a fundamental barrier to timely data analysis as file numbers get higher. Such issues arose during early development of JEX resulting in many learnings. We found that directly connected server storage (i.e., with negligible latency times) is needed for the image analysis step that provides a minimum of gigabit ethernet connection speeds and storage capacities of at least 3.5 times the size of the raw data set. Analyzed datasets can then be archived on larger and slower servers. We also found that a standard tree-based folder structure, as opposed to storage in a specialized database eliminates challenges with file access times and provides a very transparent way for users to subsequently access, compress, and transfer files after analysis using a standard file browser. With fast file transfer speeds, use of file compression slows file saving and opening, so use of uncompressed formats (e.g., TIF) and larger storage with faster read/write/transfer times is preferred. Furthermore, image analysis algorithms that can combine multiple steps into one help limit the number of intermediate image files that need to be stored, reducing read / write / transfer / processing times significantly, but can create overly specialized methods that reduce modularity for future use, requiring a well-planned design for the analysis workflow. Writing of tabular data should be performed on a line-by-line basis to disk memory to minimize RAM memory overhead. Likewise, large data tables (potentially 100’s of millions of lines long depending upon the number of cell features being measured) should be managed using memory efficient tools such as the data.table package for R.^32,33^ Also, given that identical processing steps are being performed on independent sets of information (e.g., an image set from well A1 is independent and identically formatted to an image set from well A2), multi-core processing can be readily implemented without the need to develop unique multi-threaded versions of algorithms. However, at times, limited RAM and memory intensive algorithms can limit the number of threads that can be running at once. Therefore, JEX allows the user to select the number of threads being used for each image processing operation to avoid reaching RAM memory limits that can result in data corruption or loss, while providing multi-fold reductions in processing time. Lastly, JEX was also advanced for this project to allow image processing during imaging. This allowed the ∼10 hours of raw data analysis per day of imaging to be finished within ∼15 min of completing the timelapse imaging. Given the scale and scope of these different challenges, the features of JEX for analysis of large datasets have been critical to pursuing this work successfully. As such, future implementations of similar work should be aware of such challenges and inform choices of software and hardware infrastructure. JEX provides a highly extensible open-source, Java-based framework that integrates the ImageJ2, SCIFIO, Sci-Java, and ImgLib2 libraries for developing new implementations.^34–37^

Taken together, the above technical advancements provide fundamentally new capabilities for quantifying drug response in primary cells and timelapse imaging applications at the single cell resolution.

### Detection of the mechanisms of therapeutic drug actions on both cell division and death events using a human MM cell line

With a goal to sensitively quantify cell changes in primary MM patient cells, we first developed a workflow described in Figure 1 using a test model human MM cell line, RPMI8226 cells. 15000 cells per well were seeded (typical) in collagen and treated with increasing concentrations of the clinical proteasome inhibitor bortezomib. Viable cells in each drug concentration group relative to the initial viable cells set at unity were plotted (Figure 5A). The cell number curves between hours 0 to 5 showed minimal stratification but the fraction of viable cells increased without bortezomib treatment (Bort 0 nM). In contrast, the fraction of viable cells with the drug treatment decreased over time in a dose-dependent manner. The increase in the viable cell fraction without the drug treatment relative to the start of the assay suggested that the some of the cells underwent cell division events. In the absence of bortezomib over 40 hours, the number of division events totaled ∼30% of the initial population while cell death events totaled ∼20% of the initial population (Figure 5B). The minor cell death events indicated that the health of MM cells was well maintained in the culture condition. At all bortezomib doses, cell division events were significantly and equally prevented, suggesting that a relatively low concentration of bortezomib was sufficient to completely block cell proliferation. In contrast, cell death events significantly increased in a dose- and time-dependent manner. These results demonstrated that inhibition of cell division was more sensitive to bortezomib than induction of cell death by the drug. Thus, our time-lapse live cell imaging platform was able to simultaneously quantify drug sensitivities of both MM cell division and death events over time without the use of a large number of cells and without the need to terminate cultures during imaging.

**Figure 5.**
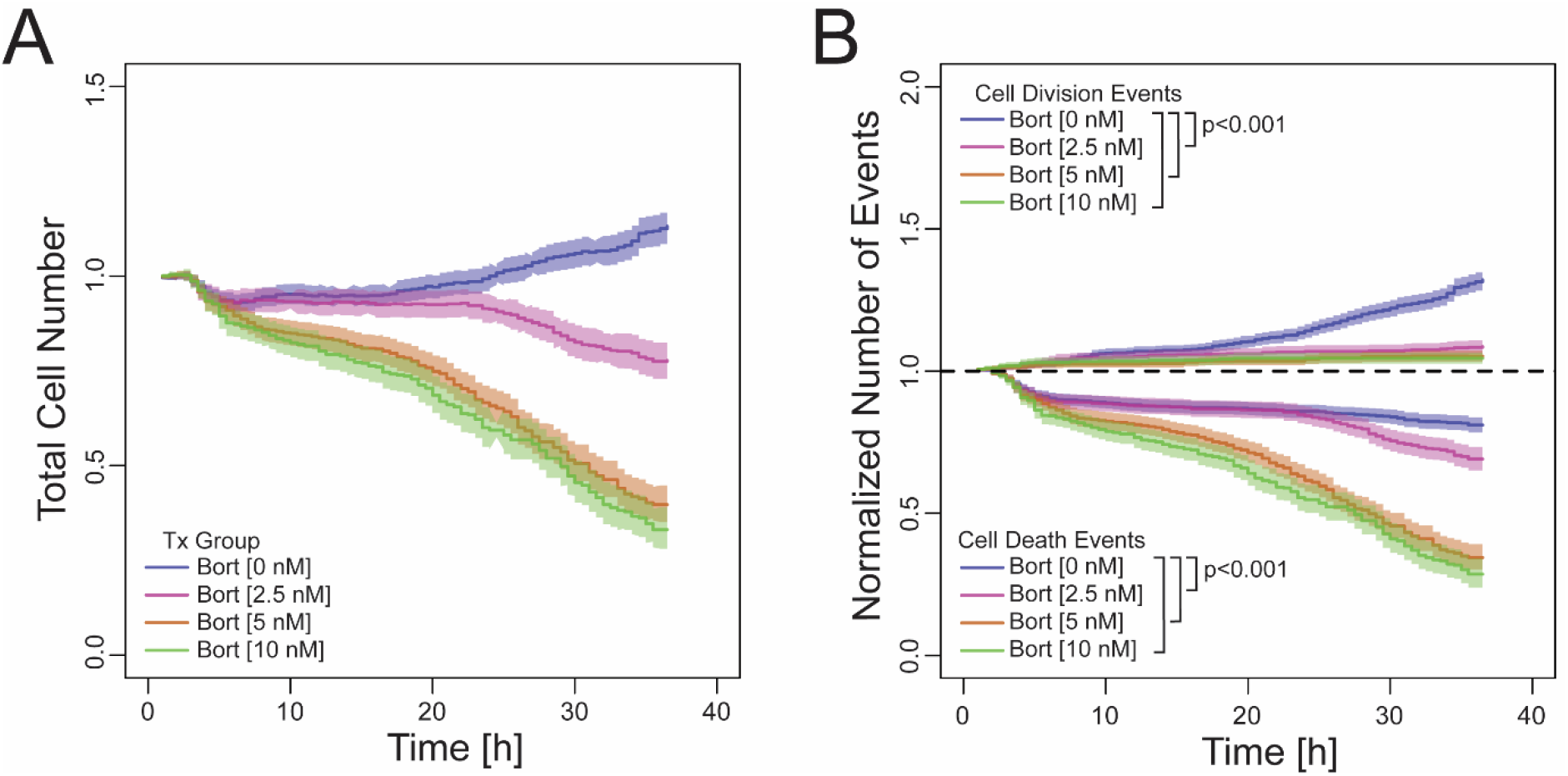
RPMI8226 cell division and death tracked over time in the presence of bortezomib. (A) Total RPMI8226 cell number over time and B) normalized number of cell division and death events after treatment with 0, 2.5, 5, or 10 nM Bort (bortezomib). Images were taken up to 72 hours. p<0.001. Single cell data from technical replicates of each condition are pooled and graphed. Total cell number is plotted on a normalized axis, normalized to the initial cell number. Number of cell death and proliferation events are plotted as the fraction of the initial cell population as well. The solid line represents the measured value for the pool and the band represents the 95% confidence interval around that measurement. The confidence interval for the total cell number (left) is calculated via propagation of uncertainty from the proliferation and death curves (right) given total cell number is initial cell number plus proliferation events, minus death events. Statistical comparisons were performed using the ‘survminer’ package in R for event / survival analyses.

To further test the capacity of our imaging tool to decipher the degree of MM cell death induced by drug treatment, RPMI8226 cells were treated with bortezomib in the presence of zVAD, a caspase inhibitor to block apoptosis, or necrostatin, a RIP1 inhibitor to block necroptosis^38,39^, and total cell number was tracked over 72 hours as above. As in Figure 5A, viable RPMI8226 cell number increased without the drugs (Figure 6A, blue). There was no difference between necrostatin + bortezomib compared to bortezomib alone (Figure 6A, green vs pink). In contrast, cell death events were significantly diminished in the presence of zVAD + bortezomib relative to bortezomib alone (Figure 6A, orange vs pink). These results are consistent with the mode of bortezomib-induced toxicity being apoptotic, not necroptotic, thus showcasing that this assay can be used to discriminate the mechanism of drug-induced cell cytotoxicity within one experiment with sensitive statistical discrimination between conditions.^40^

**Figure 6.**
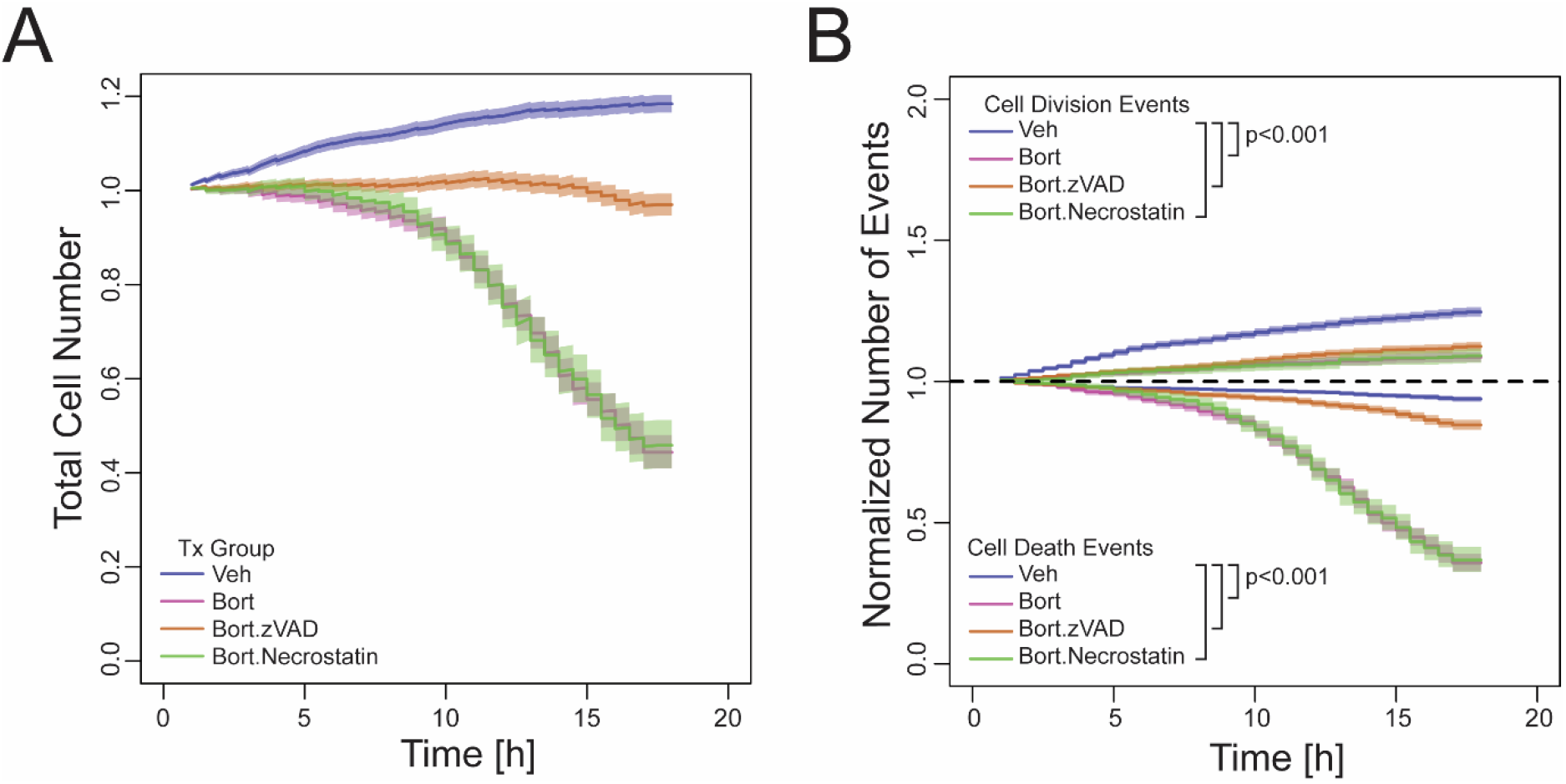
RPMI8226 cell division and death tracked over time in the presence of bortezomib, zVAD, and Necrostatin. (A) Total RPMI8226 cell number over time and B) normalized number of cell division and death events after treatment with 10 nM Bort (bortezomib) with or without 10 nM zVAD or Necrostatin. Images were taken up to 72 hours. p<0.001. Analysis was performed as described in Figure 5.

Since our assay demonstrated the ability to simultaneously quantify both cell division and death events over time, these events were separately plotted to measure the impacts of zVAD and necrostatin on bortezomib effects. Necrostatin had no significant impacts on both cell division and death events in the presence of bortezomib relative to bortezomib alone (Figure 6B, green vs pink). However, zVAD was able to significantly prevent bortezomib-induced cell death events without having any impact on bortezomib inhibition of cell division events (Figure 6B, orange vs pink). These results demonstrated that zVAD specifically prevented bortezomib-induced cell death events without affecting drug inhibition of cell division events, which is consistent with the mechanism of zVAD’s action only on apoptosis.^38^ Thus, these results showcased that our time-lapse assay is faithfully detecting drug-induced actions on both cell division and death events. Figure 5 and 6 both demonstrate that this assay can generate powerful statistics in its ability to track individual cell division or cell survival events in the presence of therapeutic drugs with a limited number of cells within a single experiment, all while providing greater insight into the mechanisms of drug actions. This helps overcome inherent difficulty in working with primary patient samples due to often limited quantities of cells with which to work and therefore severely limiting the number of replicates that can be run in parallel to obtain statistically amenable data.^19–22,41^

### Detection of the impacts of TME components on MM cell drug sensitivity

As stated previously, it is desirable to incorporate a patient’s unique TME components into assays to understand how they affect functions of cancer cells, such as drug sensitivity. We therefore incorporated patients’ bone marrow stromal cells (BMSCs) into our time-lapse RPMI8226 MM cell imaging assay along with bortezomib treatment as described in Figure 7A and in the ESI. 5 nM of bortezomib was used given it provides an appropriate degree of dose-dependent cell death (as seen in Figure 5B). The total MM cell numbers were tracked for ∼35 hours. As before, RPMI8226 cells showed increased total cell number over time (Figure 7B, green). The presence of patient BMSCs further increased the total cell number in the absence of drugs (Figure 7B, pink), suggesting that BMSCs had a pro-proliferative effect. When bortezomib was included in the assay, total viable MM cell numbers decreased without or with the presence of BMSCs (Figure 7B, orange and blue). However, it was unclear from the analysis of total viable cell numbers whether BMSCs modulated cell division or death events or both in the presence of bortezomib. By separating these cellular events, it became clear that this particular patient’s BMSCs had no statistically significant impacts on bortezomib-induced cell death events (Figure 7C, orange vs blue). Instead, the BMSCs significantly augmented MM cell division events without or with bortezomib (Figure 7C, upper curves). The results establish that a second cell type can be effectively incorporated into our time-lapse cell imaging assay without compromising the health of MM cells and to reveal the cellular mechanism of the impact of stromal elements on tumor cell fate in the presence of a therapeutic drug.

**Figure 7.**
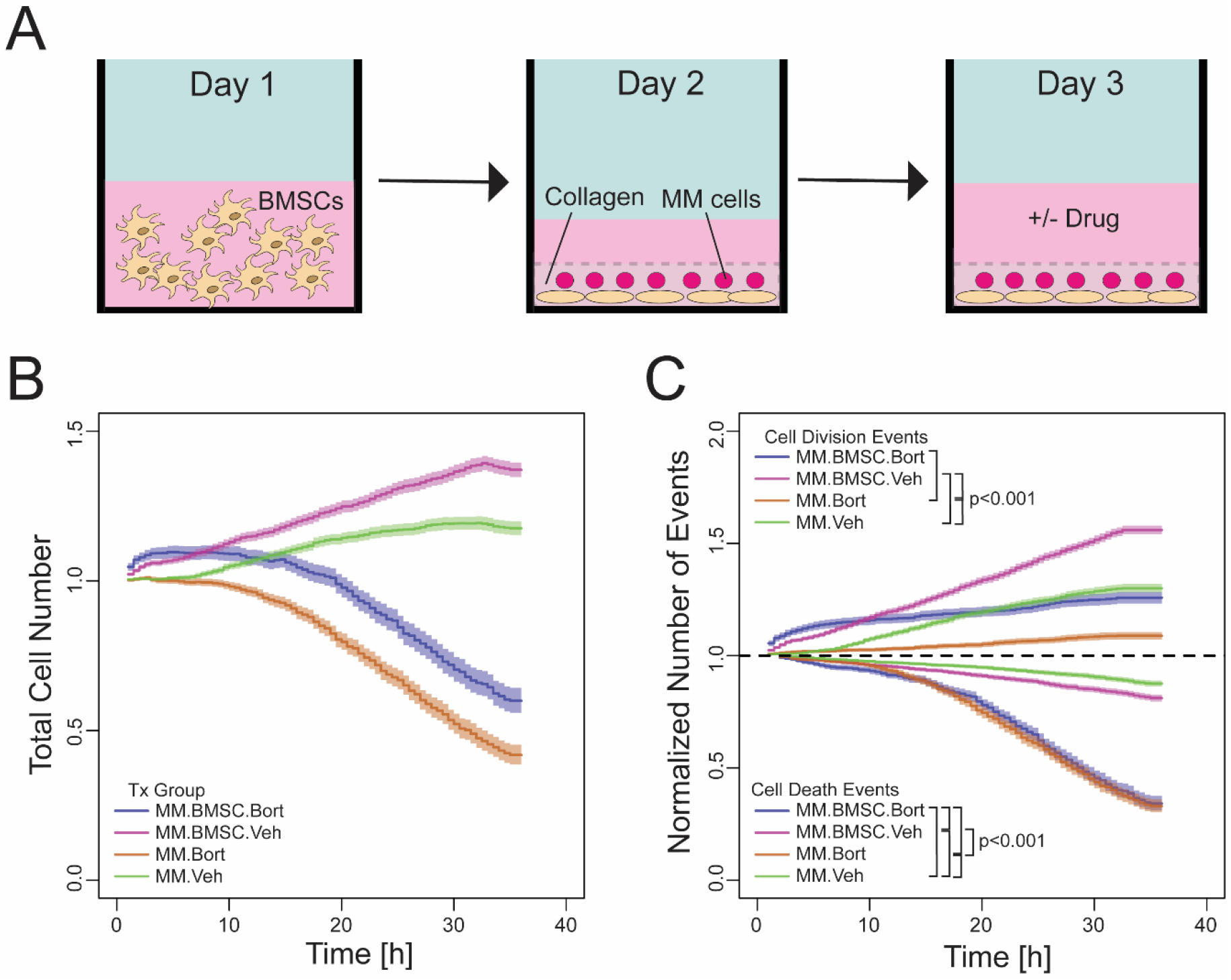
Multiple myeloma and bone marrow stromal cells time-lapse workflow (above). BMSCs are plated overnight in a 96-well glass bottom dish before myeloma cells are laid on top with collagen the following day. The two cell types are allowed to interact for 24 hours before drugs and media are dispensed on top of the co-culture. RPMI8226 cell division and death tracked over time in the presence of BMSCs and bortezomib (below). Total RPMI8226 cell number over time and (C) normalized number of cell division and death events after treatment with 5 nM Bort (bortezomib) with or without BMSCs. Images were taken up to 72 hours. p<0.001. Analysis was performed as described in Figure 5.

### Detection of primary MM cell sensitivity to multiple therapeutic drugs in the presence of patient-matched TME components

Our previous studies have found that BMSCs established from different MM patients display heterogeneity in their functional impacts on MM cells, such as cell signaling^42^and drug responses^43^, thus emphasizing the potential need to incorporate patient-matched TME components in *ex vivo* assays for better prediction of clinical outcomes.^22^ Thus, to determine the utility of our time-lapse assay in analyzing drug sensitivities of primary patient MM cells in the presence of their own TME components, we next separated the CD138^+^ MM cell fraction from the CD138^-^mononuclear cell fraction (containing all other cell types) from patient bone marrow aspirates. We also isolated the plasma fraction from the bone marrow aspirates which includes other soluble TME components. Primary CD138^+^cells were embedded within collagen matrix and bone marrow plasma and an equal number of the CD138^-^cell fraction was overlaid over the collagen gel to match the patient’s own soluble and cellular TME components. These cells were then treated with established myeloma drugs, bortezomib, dexamethasone, carfilzomib, Selinexor, or the triple combination of carfilzomib, dexamethasone, and Selinexor. As in RPMI8226 cell line, the total number of primary CD138^+^ MM cells increased over the course of 38 hours (Figure 8A, purple), indicating that the health of these primary MM cells is maintained, and cell division events are also supported in this culture condition. Dexamethasone (pink) and Selinexor (light green) caused no and small reduction, respectively, on total cell number. In contrast, bortezomib (purple) and carfilzomib (orange), both being proteasome inhibitors, caused similar reductions in total cell number. A combination of carfilzomib, Selinexor and dexamethasone (green) had the greatest reduction in total MM cell number relative to vehicle alone (blue). By separating cell division and death events, it became clear that only the triple drug combination had significant inhibition of CD138^+^ MM cell division over no drug or other individual drug conditions (Figure 8B, green vs all others). In contrast, Selinexor, bortezomib, and carfilzomib all reduced the cell survival events relative to vehicle or dexamethasone alone conditions. Like in the cell division events, the triple combination had the greatest reduction in cell survival events (Figure 8B, green). Thus, the ability to simultaneously quantify the cell division and death events from a limited number of primary CD138^+^ MM cells in the presence of patient’s own TME components under different drug treatment conditions enabled the elucidation of cellular mechanisms of individual and combined drugs actions. Figure 8 demonstrates that this tool can measure a functional and phenotypic readout for cells in the presence of patient-matched TME components without and with various therapeutic drugs. These technical advancements provide unique alternatives to endpoint survival assays of a single readout, such as viability, colorimetric, and clonogenic assays, and to assays which depend on passaging primary cells over time, possibly changing the fractions of their genetic variants.^44–46^

**Figure 8.**
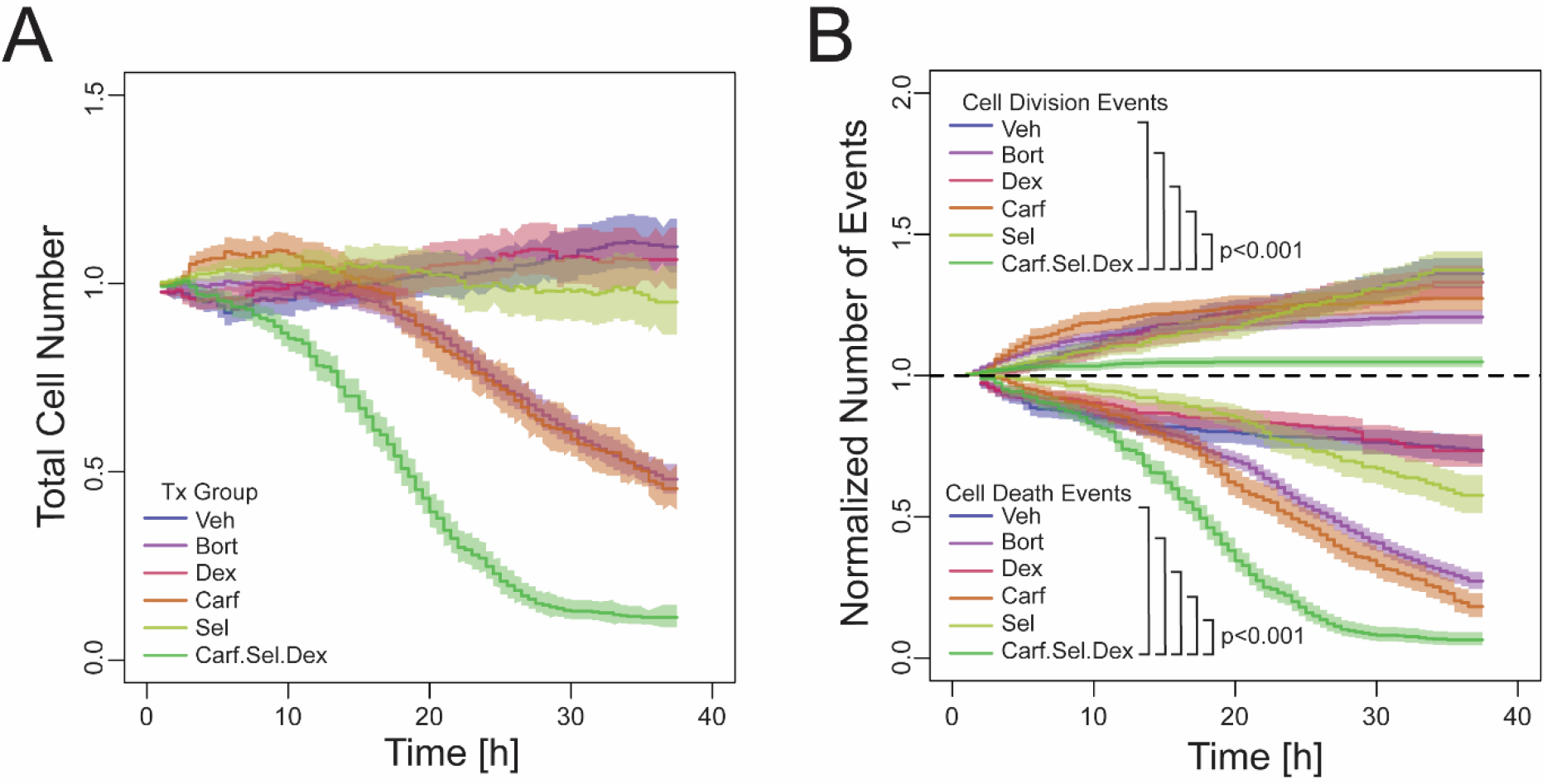
Primary myeloma cell division and death tracked over time in the presence of patient-specific plasma and other populations and various myeloma treatments. (A) Total CD138+ cell number over time and B) normalized number of cell division and death events after treatment with 10 nM Bort (bortezomib), 100 nM Dex (dexamethasone), 10 nM Carf (carfilxomib), 10 nM Sel (Selinexor) alone or in combination. Images were taken up to 72 hours. p<0.001. Analysis was performed as described in Figure 5.

Our assay also resolves a notable hurdle that other assays face in maintaining primary MM tumor cells *ex vivo*. Primary MM cells are often prone to stress and undergo apoptosis once isolated from bone marrow aspirate without the addition of exogenous cytokines, such as IL-6, or BMSCs that produce such cytokines.^14,16^ The loss of primary MM cell viability in the absence of any drugs was inherent in our previous µC3 assay system and was also considered a weakness of the system.^22^ The current assay can factor in TME cues such as hypoxia, pH gradients, and organization and stiffness of the extracellular matrix using 3D collagen. The increase in total numbers of primary MM cells with limited cell death events in the absence of any drug over time shown in Figure 8 showed that the health of primary MM cells was well maintained under the culture conditions. Moreover, the utilization of patient-matched TME components, such as bone marrow plasma, the CD138^-^cell fraction or a specific factor or cell type within these TME components, gave patient-specific insights into drugs’ impacts on cell death and division events. Thus, this time-lapse 3D assay platform provides a new avenue to develop it as a sensitive diagnostic assay which can model the tumor cell and environmental heterogeneity to test permutations of drugs and to eventually predict individual patient’s therapeutic response.

### Future Perspectives

Beyond improving endpoint survival readouts, this time-lapse imaging assay provides an expansive opportunity to combine the study of the phenotypic heterogeneity of individual cells with single-cell omics approaches. Current efforts to bridge the connection between digital image analyses and single-cell omics analyses often involve precisely isolating single cells in a heterogenous population through microfluidics-based techniques after imaging. Some examples include encapsulating single cells within a microwell or a water droplet, or dielectrophoretically sorting them.^47–49^ In our system, it is possible to merge this time-lapse imaging technique aided by the software JEX with an inexpensive semiautomated single-cell aspirator (SASCA) system that can enrich rare cell populations by immobilizing cells from impure populations into microwells that will then undergo whole transcriptome and genome analysis via scRNA.^52,53^ Thus, merging our time-lapse assay and the SASCA would allow us to observe dynamic phenotypic changes within each cell continuously rather than collecting individual timepoints, which would only provide a functional endpoint view of cellular behavior and heterogeneity. Although not leveraged here, one should also note that hundreds of different metrics exist for quantifying cell shape, intensity distribution, and context (e.g., types of adjacent cells) and can be integrated with machine learning techniques to further advance quantification of cell phenotypes and prediction or discrimination of responses in future studies. The tools presented here provide a foundation for developing such approaches and complementing them with single-cell “omics” endpoints to further improve our understanding of primary tumor cell heterogeneity in a patient-individualized fashion.

## Supporting information

Paper ESI

## Acknowledgments

This work was supported by Cancer Biology Training Grant T32 CA009135 NRSA Award (to CM), Collaborative Health Sciences Program Award #AAB7473 under Wisconsin Partnership Program (to SM and JW), NIH/NCI R01 CA155192 and R01 CA251595 (to SM), A RIDE Scholar Award (to SM), and a pilot fund from the UW Carbone Cancer Center Grant P30 CA014520 (to SM). The authors thank David Page, Sean Mcilwain, Irene Ong for helpful data analysis discussions; Curtis Rueden and the LOCI team for their efforts to develop ImageJ2 libraries that enabled integration of image analysis capabilities into these studies; Dr. Zuo from the Nanjing University of Science and Technology for graciously providing us with the Matlab code referenced in the study; the Department of Hematology Clinical Trials Staff for help with IRB protocols and sample acquisition; all the members of the Miyamoto lab for fruitful discussions and suggestions, and all the MM patients for donating study samples.

## Materials and Methods

The ‘Key Technical Components of the Assay’ section provides description and discussion of the fundamental methods and approaches used to perform this study. Additional technical details are contained in the ESI, including cell culture methods, drug treatments, mathematical descriptions of algorithms, and links to the open-source code repositories.

## References

1. Sasagawa, Y. et al. Quartz-Seq: a highly reproducible and sensitive single-cell RNA sequencing method, reveals non-genetic gene-expression heterogeneity. Genome Biology 14, 3097 (2013).

2. Goldman, S. L. et al. The Impact of Heterogeneity on Single-Cell Sequencing. Frontiers in Genetics 10, 8 (2019).

3. Choi, Y. H. & Kim, J. K. Dissecting Cellular Heterogeneity Using Single-Cell RNA Sequencing. Mol Cells 42, 189–199 (2019).

4. Hou, Y. et al. Single-cell triple omics sequencing reveals genetic, epigenetic, and transcriptomic heterogeneity in hepatocellular carcinomas. Cell Res 26, 304–319 (2016).

5. Nam, A. S., Chaligne, R. & Landau, D. A. Integrating genetic and non-genetic determinants of cancer evolution by single-cell multi-omics. Nat Rev Genet 22, 3–18 (2021).

6. Mossner, M., Baker, A.-M. C. & Graham, T. A. The role of single-cell sequencing in studying tumour evolution. Fac Rev 10, 49 (2021).

7. Henke, E., Nandigama, R. & Ergün, S. Extracellular Matrix in the Tumor Microenvironment and Its Impact on Cancer Therapy. Frontiers in Molecular Biosciences 6, 160 (2020).

8. Winkler, J., Abisoye-Ogunniyan, A., Metcalf, K. J. & Werb, Z. Concepts of extracellular matrix remodelling in tumour progression and metastasis. Nat Commun 11, 5120 (2020).

9. Ghosh, N., Ye, X., Ferguson, A., Huff, C. A. & Borrello, I. Bortezomib and thalidomide, a steroid free regimen in newly diagnosed patients with multiple myeloma. Br J Haematol 152, 593–599 (2011).

10. Chen, D., Frezza, M., Schmitt, S. & Dou, J. K. and Q. P. Bortezomib as the First Proteasome Inhibitor Anticancer Drug: Current Status and Future Perspectives. Current Cancer Drug Targets http://www.eurekaselect.com/73304/article (2011).

11. Sinha, S. et al. Impact of Dexamethasone Responsiveness on Long Term Outcome in Patients with Newly Diagnosed Multiple Myeloma. Br J Haematol 148, 853–858 (2010).

12. Mazumder, A. & Jagannath, S. Thalidomide and lenalidomide in multiple myeloma. Best Practice & Research Clinical Haematology 19, 769–780 (2006).

13. Myeloma - Cancer Stat Facts. SEER https://seer.cancer.gov/statfacts/html/mulmy.html.

14. Hideshima, T., Mitsiades, C., Tonon, G., Richardson, P. G. & Anderson, K. C. Understanding multiple myeloma pathogenesis in the bone marrow to identify new therapeutic targets. Nature Reviews Cancer 7, 585–598 (2007).

15. Bar-Natan, M. et al. Bone marrow stroma protects myeloma cells from cytotoxic damage via induction of the oncoprotein MUC1. Br J Haematol 176, 929–938 (2017).

16. Chatterjee, M. et al. In the presence of bone marrow stromal cells human multiple myeloma cells become independent of the IL-6/gp130/STAT3 pathway. Blood 100, 3311–3318 (2002).

17. Chesi, M. et al. AID-dependent activation of a MYC transgene induces multiple myeloma in a conditional mouse model of post-germinal center malignancies. Cancer Cell 13, 167–180 (2008).

18. Rajagopalan, A. et al. Mice Expressing MYC and NrasQ61R in Germinal Center B Cells Develop Highly Aggressive Multiple Myeloma. Blood 132, 1006–1006 (2018).

19. Zhou, Y. et al. Growth Control of Multiple Myeloma Cells through Inhibition of Glycogen Synthase Kinase-3. Leuk Lymphoma 49, 1945–1953 (2008).

20. Jakubikova, J. et al. Lenalidomide targets clonogenic side population in multiple myeloma: pathophysiologic and clinical implications. Blood 117, 4409–4419 (2011).

21. Walker, Z. J. et al. Measurement of ex vivo resistance to proteasome inhibitors, IMiDs, and daratumumab during multiple myeloma progression. Blood Adv 4, 1628–1639 (2020).

22. Pak, C. et al. MicroC3: an ex vivo microfluidic cis-coculture assay to test chemosensitivity and resistance of patient multiple myeloma cells. Integr. Biol. 7, 643–654 (2015).

23. Khin, Z. P. et al. A Preclinical Assay for Chemosensitivity in Multiple Myeloma. Cancer Res 74, 56–67 (2014).

24. Mir, M., Bhaduri, B., Wang, R., Zhu, R. & Popescu, G. Chapter 3 - Quantitative Phase Imaging. in Progress in Optics (ed. Wolf, E.) vol. 57 133–217 (Elsevier, 2012).

25. Warrick, J. W. & Berthier, E. JEX [Software]. (2021).

26. Zuo, C., Chen, Q., Li, H., Qu, W. & Asundi, A. Boundary-artifact-free phase retrieval with the transport of intensity equation II: applications to microlens characterization. Opt Express 22, 18310–18324 (2014).

27. Volkov, V. V., Zhu, Y. & De Graef, M. A new symmetrized solution for phase retrieval using the transport of intensity equation. Micron 33, 411–416 (2002).

28. Zhang, X. & Kiechle, F. L. Hoechst 33342-induced apoptosis in BC3H-1 myocytes. Ann Clin Lab Sci 27, 260–275 (1997).

29. Zhang, X., Chen, J., Davis, B. & Kiechle, F. Hoechst 33342 Induces Apoptosis in HL-60 Cells and Inhibits Topoisomerase I In Vivo. Archives of Pathology & Laboratory Medicine 123, 921–927 (1999).

30. Klak, M. et al. Irradiation with 365 nm and 405 nm wavelength shows differences in DNA damage of swine pancreatic islets. PLOS ONE 15, e0235052 (2020).

31. Purschke, M., Rubio, N., Held, K. D. & Redmond, R. W. Phototoxicity of Hoechst 33342 in time-lapse fluorescence microscopy. Photochem. Photobiol. Sci. 9, 1634– 1639 (2010).

32. RStudio Team and Others. RStudio: Integrated Development for R. https://rstudio.com/ (2021).

33. R Core Team. R: a language and environment for statistical computing. https://www.R-project.org/ (2021).

34. Rueden, C. T. et al. ImageJ2: ImageJ for the next generation of scientific image data. BMC Bioinformatics 18, 529 (2017).

35. Pietzsch, T., Preibisch, S., Tomancák, P. & Saalfeld, S. ImgLib2--generic image processing in Java. Bioinformatics 28, 3009–3011 (2012).

36. Hiner, M. C., Rueden, C. T. & Eliceiri, K. W. SCIFIO: an extensible framework to support scientific image formats. BMC Bioinformatics 17, 521 (2016).

37. Rueden, C. SciJava-common -- a generic Java framework for science. SCIJAVA-COMMON PLUGINS, CONTEXTS, UTILITIES AND MORE https://scijava.org/scijava-common/scijava-common.html#/1.

38. Wu, Y.-T. et al. zVAD-induced necroptosis in L929 cells depends on autocrine production of TNFα mediated by the PKC–MAPKs–AP-1 pathway. Cell Death Differ 18, 26–37 (2011).

39. Vandenabeele, P., Grootjans, S., Callewaert, N. & Takahashi, N. Necrostatin-1 blocks both RIPK1 and IDO: consequences for the study of cell death in experimental disease models. Cell Death Differ 20, 185–187 (2013).

40. Park, J. et al. Establishment and characterization of bortezomib-resistant U266 cell line: Constitutive activation of NF-κB-mediated cell signals and/or alterations of ubiquitylation-related genes reduce bortezomib-induced apoptosis. BMB Rep 47, 274–279 (2014).

41. Warrick, J. W., Young, E. W. K., Schmuck, E. G., Saupe, K. W. & Beebe, D. J. High-content adhesion assay to address limited cell samples. Integrative Biology 5, 720– 727 (2013).

42. Markovina, S. et al. Bortezomib-Resistant Nuclear Factor-κB Activity in Multiple Myeloma Cells. Mol Cancer Res 6, 1356–1364 (2008).

43. Markovina, S. et al. Bone marrow stromal cells from multiple myeloma patients uniquely induce bortezomib resistant NF-κB activity in myeloma cells. Molecular Cancer 9, 176 (2010).

44. Franken, N. A. P., Rodermond, H. M., Stap, J., Haveman, J. & van Bree, C. Clonogenic assay of cells in vitro. Nat Protoc 1, 2315–2319 (2006).

45. Riss, T. L. et al./person-group>. Cell Viability Assays. in Assay Guidance Manual (eds. Markossian, S. et al.) (Eli Lilly & Company and the National Center for Advancing Translational Sciences, 2004).

46. Januszyk, M. et al. Evaluating the Effect of Cell Culture on Gene Expression in Primary Tissue Samples Using Microfluidic-Based Single Cell Transcriptional Analysis. Microarrays (Basel) 4, 540–550 (2015).

47. Potter, N. E. et al. Single-cell mutational profiling and clonal phylogeny in cancer. Genome Res 23, 2115–2125 (2013).

48. Joensson, H. N. & Andersson Svahn, H. Droplet Microfluidics—A Tool for Single-Cell Analysis. Angewandte Chemie International Edition 51, 12176–12192 (2012).

49. Peeters, D. J. E. et al. Semiautomated isolation and molecular characterisation of single or highly purified tumour cells from CellSearch enriched blood samples using dielectrophoretic cell sorting. Br J Cancer 108, 1358–1367 (2013).

50. Kamal, M. et al. PIC&RUN: An integrated assay for the detection and retrieval of single viable circulating tumor cells. Sci Rep 9, 17470 (2019).

51. Parker, S. G. et al. A photoelectrochemical platform for the capture and release of rare single cells. Nat Commun 9, 2288 (2018).

52. Bai, D., Peng, J. & Yi, C. Advances in single-cell multi-omics profiling. RSC Chemical Biology 2, 441–449 (2021).

53. Tokar, J. J. et al. Pairing Microwell Arrays with an Affordable, Semiautomated Single-Cell Aspirator for the Interrogation of Circulating Tumor Cell Heterogeneity. SLAS TECHNOLOGY: Translating Life Sciences Innovation 25, 162–176 (2020).

54. Tinevez, J.-Y. et al. TrackMate: An open and extensible platform for single-particle tracking. Methods 115, 80–90 (2017).

55. Chenouard, N. Objective comparison of particle tracking methods | Nature Methods. https://www.nature.com/articles/nmeth.2808.

