## Supplementary material for "Timelapse viability assay to detect division and death of primary multiple myeloma cells in response to drug treatments with single cell resolution": Paper ESI

**Supplemental Information**

*Cell lines, chemicals, and drugs*

RPMI8226 and 293T cell lines were purchased from American Type Culture Collection (ATCC). RPMI8226 cells were cultured in RPMI1640 media containing 10% FBS, 2% GlutaMAX (Gibco), and 1% penicillin/streptomycin, and grown in 37 °C/5% $CO_2$. 293T cells were cultured in Dulbecco's Modified Eagle Medium (DMEM) containing 10% FBS and 1% penicillin/streptomycin. Bortezomib (S1013, Selleckchem), carfilzomib (S2853, Selleckchem), and zVAD-fmk (550377, BD Biosciences) were dissolved in DMSO at 10 mM stock solutions. necrostatin (AG-CR1-2900-M005, Adipogen) was dissolved in DMSO into 100 mM stock solution. Dexamethasone (D4902, Sigma Aldrich) was dissolved in DMSO at 100 mM stock solution. Selinexor (KPT-330, Karyopharm) was DMSO into 10mM stock solution. Hoechst 33258 (H3569, Thermo Fisher) was dissolved in water at 16 mM stock solution. Vehicle controls were treated with DMSO alone.

*Primary cell isolation and culture*

Primary human MM and BMSCs were obtained from fresh whole bone marrow aspirates with informed consent in accordance to the University of Wisconsin Institutional Review Board requirements (HO07403) at the University of Wisconsin Hospital and Clinics. Pt. 829 (Figure 7) was derived from a male patient who was of age 70 and relapsed on a combination of Velcade (bortezomib), Revlimid (lenalidomide), dexamethasone, and elotuzamab with 30% plasma cells on biopsy. Pt. 849 (Figure 8) was derived from a newly diagnosed female patient who was of age 50 with 63% plasma cells on biopsy. Bone marrow aspirates were spun at 400 x g for 5 min at room temperature in order to collect the BM plasma fraction for further analysis (stored in -80°C). The mononuclear fraction was then positively sorted for myeloma plasma cells using $CD138^+$ magnetic MACS beads (Miltenyi Biotec) to $\geq$90% purity as described previously.[1] $CD138^-$ fractions were cultured in OptiMEM media containing 10% FBS, 2% GlutaMAX, 1% NEAA (Gibco), and 1% penicillin/streptomycin and incubated at 37°C in a 5% $CO_2$ incubator. The mononuclear fraction was then positively sorted for myeloma plasma cells using $CD138^+$ magnetic MACS beads (Miltenyi Biotec) to 90% purity as described previously.[1] $CD138^-$ fractions were cultured in OptiMEM media containing 10% FBS, 2% GlutaMAX, 1% NEAA (Gibco), and 1% penicillin/streptomycin and incubated at 37 °C in a 5% $CO_2$ incubator. Nonadherent cells were removed 24 h after plating to allow BMSCs to grow out. Adherent BMSCs were passaged 3-4 times and cryopreserved as previously described.[1]

*Mono and co-culture assay*

For the monoculture, RPMI8226 cells or primary $CD138^+$ cells were plated at 15,000 cells per well in a 96-well plate in OptiMEM whole media (containing 10% FBS, 2% GlutaMAX, 1% NEAA (Gibco), and 1% penicillin/streptomycin), mixed with a collagen gel (containing 10% 10X PBS, 8.55% tissue culture-grade $H_2O$, 1.45% 1N NaOH, and 80% PureCol (cat # 5005)), dispensed within designated well, spun down at 250xg at room temperature, and then incubated for 45 minutes at 37°C to allow gel to solidify. For the BMSC co-culture, primary BMSCs were trypsinized and plated at

10,000 cells per well in OptiMEM whole media to adhere overnight. After 24 hours, RPMI8226 cells in collagen were plated on top as described above. For the CD138$^+$ co-culture, CD138$^+$ cells were isolated and prepared in collagen, as stated above, and CD138$^-$ cells and plasma from the corresponding patient sample were dispensed on top of the collagen-cell mixture after it solidified.

*Drug treatments*

RPMI8226 cells were treated with 5 or 10nM bortezomib with or without 10 μM zVAD-fmk or 10 μM nectrostatin. CD138$^+$ cells were treated with 10 nM carfilzomib, 100 nM dexamethasone, and 10 nM Selinexor. Designated drug treatment and 1:500 of 200 μg/mL Hoechst stain were added on top of BMSC and RPMI collagen-containing wells. Wells contain a total of 200uL of collagen (+ cells) and media.

*Live cell imaging 3D (co-)culture device and microscopy*

TI-SH-U incubation stage (>95% relative humidity, 5 $\pm$ 1 % $CO_2$, and 37 $\pm$ 1°C) warmed up prior to drug and 1:500 of 200 μg/mL Hoechst stain. After wells were treated with either vehicle or drug, 96-well plate was set inside the humidity chamber, Nikon TI-S-EJOY microscope was programmed to the desired well locations (4 rows, 4 columns, 3X3 image montage within each well), MM cells were put into focus, acquisition parameters were set (mean background BF intensity 30,000 units out of 65535 at 10 ms exposure, 395 nm fluorescence exposure time of 100 ms to maximize cell health with peak live cell nuclear intensity 10-15% above background fluorescence levels of ~1300 units), and imaging was started. Cells were imaged at 30-minute intervals for at least 35 hours. A total of 3 images were taken at each location within the 3X3 montage. The fluorescent image was acquired at the z=0 focal plane while a brightfield image was taken at both z = +3 and -3 µm for input in the TIE.

*QPI algorithm*

Java algorithm is derived from MATLAB code generously provided by Dr. Chao Zuo at the Smart Computational Imaging Laboratory (SCILab), Nanjing University of Science and Technology. The mathematical basis for the algorithm is described in detail in previous publications.[2] Briefly, the TIE equation for small deviations in z is described by Equation 1 for non-uniform incident intensities and simplified to a Poisson-form in Equation 2 when illumination is constant across the field of view. *I* is the Intensity field while $I_0$ is the incident light intensity field. For our microscope, incident illumination was highly uniform across the 20x field of view for brightfield and Equation 2 was used.

$$\phi = -\frac{2\pi}{\lambda}\nabla^{-2}\left[\nabla\cdot\left(\frac{1}{I}\nabla\nabla^{-2}\frac{\partial I}{\partial z}\right)\right],\ k = \frac{2\pi}{\lambda} \tag{1}$$

$$\phi = -\frac{k}{I_0}\nabla^{-2}\frac{\partial I}{\partial z} \tag{2}$$

We then used the Fast-Fourier-Transform ($\mathcal{F}$) method in Equation 3 with even symmetrization on image data, *u(x, y)*, to calculate Equation 2.[3]

$$\nabla^{-2}u(x,y) = \mathcal{F}^{-1}\left[\frac{\mathcal{F}[u(x,y)]}{|q|^2}\right] \tag{3}$$

Division by $|q|^2$ in Equation 3 creates a high-pass filter where the characteristic cutoff frequency is determined by the value of q. In this work, q was set to 32/100 where 100 is the feature size cutoff in pixels of the filter, retaining features below this cutoff. $\partial I/\partial z$ was calculated from two images spaced 6 µm apart along the z-axis. The dominant wavelength of light, λ, was assumed to be 600 nm for our LED brightfield illumination. The camera pixel size was 6.45 µm making each image pixel at 20x magnification 0.3225 µm.

*Weighted-mean background subtraction*

As the name implies, weighted-mean background subtraction subtracts the local (as determined by the kernel being used) weighted-mean from the image to perform background subtraction. This is very similar to standard mean background subtraction, but the weight given to each pixel in the mean is determined by (i) a weighting "mask", and (ii) the weights given to each location in the filter kernel (e.g., uniform weighted kernel or Gaussian weighted kernel). "Mask" is put into quotes here as this mask can have a continuous range or be binary. The weighting mask provides a way for users to specify user-specific algorithms for identifying pixels that are more or less likely to be background pixels while the kernel filter kernel functions as a way to apply local spatial weighting to those weights (e.g., weights near the center of the kernel can have more influence than weights near the kernel boundary). This method allows for an infinite number of possible weighting schemes for different applications in a logical and computationally efficient manner. While the relatively simple concept of a weighted mean is not new to math and computation, we have not expressly encountered examples in the literature or implemented in software libraries that describe such a method for image background subtraction and, thus, have described it here. The primary challenge is to come up with a fast and effective way to calculate background mask weights that give as much weight as possible to background pixels (e.g., 1) and as little relative weight as possible to foreground pixels (i.e., 0).

The primary weighting mask used here is calculated based upon the pixel-wise variance (or more precisely the pixel-wise standard deviation) in the original image. A standard deviation filter with a kernel size that is relatively small compared to objects of interest helps preserve image detail while being large enough to observe significant changes in intensity across object boundaries and features. In this case, a typical cell is on the order of 30 pixels in diameter. Thus, a kernel radius of ~2 pixels is appropriate and can be varied empirically to observe its impact on detail retention. Here a value of 2 was used for fluorescent images and a value of 4 was used for phase images. In the background, the value of the standard deviation is typically low and relatively constant while it is higher and more variable at foreground pixels. Therefore, a histogram of the standard deviations of the image should have a peak associated with the background pixels. In practice, what is seen is that the background pixels generally show the lowest standard deviations and have a generally Gaussian-like profile with an extended tail to the right. Ideally, we would like to give pixels within this peak the highest weights and values outside this peak the lowest weights. Therefore, we simply measured the width of this peak to aid in the subsequent weighting process by measuring the width of the left half of the peak, starting from the histogram mode ($h_{mode}$) (i.e., the top of the peak that estimates the median background standard deviation), down to the 0.5% percentile value ($h_{min}$). The histogram for this procedure was made on the lower 50% of standard deviation values to increase histogram resolution and used Equation 4 to determine the number of histogram, $n_{bins}$, where $n_{pixels}$ is the number of pixels in the image, $\text{bins}_{\text{min}}$ (set to 10) and $bins_{max}$ (set to 250) are the minimum and maximum allowed number of bins, respectively.

$$n_{bins} = \max\left(bins_{min}, \frac{\sqrt{n_{pixels}}\ \mathrm{bins}_{max}}{bins_{max} + \sqrt{n_{pixels}}}\right) \tag{4}$$

We then determine standard deviation values associated with the center of the first histogram bin $h_{min}$, and the center of the mode bin of the histogram. Once $h_{min}$ and $h_{mode}$ were determined, the standard deviation values from the image are transformed into weights ($w_f$) using the following two-step algorithm.

$$w_i(m,n) = \left\{ \begin{array}{ll} \delta & \text{for } x(m,n) = 0 \\ \frac{1}{1+\left(\frac{|sd(m,n)-h_{\text{mode}}|}{h_{\text{mode}}-h_{\text{min}}}\right)^p} & \text{for } x(m,n) > 0 \end{array} \right\} \tag{5}$$

$$w_f(m,n) = \left\{ \begin{array}{lr} \delta & \text{for } w_i(m,n) < \delta \\ \delta & \text{for !Double.isFinite}(w_i(m,n)) \\ 1 & \text{for } w_i(m,n) > 1 \\ w_i(m,n) & \text{otherwise} \end{array} \right\} \tag{6}$$

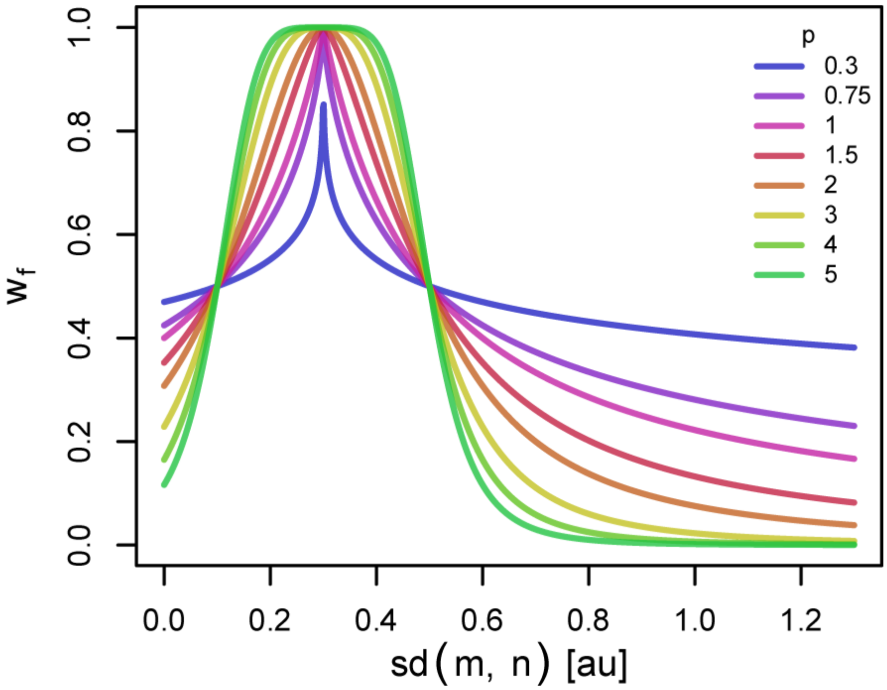


**Supp. Figure 1** – Plot of weighting function as a function of *sd(m,n)* for $\delta=10^{-7}$, $h_{mode}=0.3$, $h_{min}=0.1$, and different scaling factor values, *p*. The weighting function is a symmetric around the value of $h_{mode}$, with a maximum value of 1 at $sd(m,n)=h_{mode}$. The function reduces to 0.5 at $sd(m,n)=h_{mode}\pm(h_{mode}-h_{min})$ and rapidly approaches 0 as $sd(m,n)\rightarrow\pm\infty$. Different values of *p* change how quickly weights drop off above and below $h_{mode}$ thereby controlling how sharply weighted background pixels are in the weighted mean calculation. Also notice that the closer the values of $h_{mode}$ *and* $h_{min}$ are to one another, the tighter the weighting scheme is focused around $h_{mode}$. Effective values of p are generally between 0.3 and 5 with a value of 2 being used for our datasets.

It is possible to use the weighting mask as calculated from Eq 6; however, one can also threshold this weighting mask to create a binary weighting mask or even threshold and adjust it using binary filters such as an opening filter (erosion followed by dilation) to remove spurious points in the binary mask. One can also use other methods to identify background pixels and use an OR operation to merge them into a combined estimate of where background pixels lie. The binary mask can also be converted back to a gray-scale mask by blurring features to soften the weight estimates at object boundaries. In the work presented here, the weighting mask from Equation 6 was directly used for *fluorescence* image background correction.

In contrast, *phase* image correction relied on two weighting masks. The first was the weighting mask of Equation 6, thresholded at a value of 0.5, followed by an opening filter (erosion followed by dilation) with a radius of 2 times the radius of the variance filter. We merged the blurred variance weighted mask with the binary mask created from the lower 0.5% percentile pixels of the rolling-ball filtered phase image (ImageJ rolling-ball background subtraction filter set to the paraboloid method with a filter radius of 5 pixels), using an OR operation to "add" the 1-weighted pixels of the rolling ball mask to the variance weighting mask. The rolling-ball filter mask helps to identify extreme local minima in the phase image (typically between tightly packed cells) that should be considered background but have standard deviations that are too high to show up in the variance weighted mask. In contrast, the variance weighted mask does better at cell boundaries and in open areas between cells. Combining them retains the strengths of both. The merged weighting masks were then blurred with a radial decay blurring filter to soften weighting at cell boundaries to create the final weighting mask for the phase images. The final weights, $kw_f$, of the radial decay filter kernel were defined by Equations 7 and 8, where $x$ and $y$ are the pixel locations within the kernel relative to its center; the kernel radius, $r$, was set to 5 pixels; and the power, $p$, was set to 3.5.

$$kw_i(x,y) = \frac{1}{1+\left(\frac{\sqrt{x^2+y^2}}{r}\right)^p} \tag{7}$$

$$kw_f(x,y) = \frac{kw_i(x,y)}{\sum kw_i(x,y)} \tag{8}$$

Using the weighting masks described above for fluorescent and phase images, Equation 9 is used to calculate the estimated image background ($I_{bg}$), where $I_i$ is the initial or original image, $I_w$ is the weighting mask, and $f_k()$ is a Gaussian mean/blur filter with a radius of $r$ = 100 pixels that represents the standard deviation of the Gaussian profile used to generate the weights of the kernel. Here multiplication and division are simple pixel-wise operations and not matrix algebra.

$$I_{bg} = \frac{f_k\left(I_0 \times I_w\right)}{f_k\left(I_w\right)} \tag{9}$$

At this point, one must choose how that estimated background will be used to correct the image. One may choose to subtract the background from the original signal (i.e., $I_{final} = I_0 - I_{bg}$) or potentially divide the original signal by the background (i.e., $I_{final} = I_0/I_{bg}$). Division by background often makes sense for fluorescent applications where backgrounds and signals are often proportional to treatment concentrations and exposure times. However, instead of simply dividing by the background, which scales the image such that the mean background intensity is a value of 1, we first measure the mean background intensity, $I_{bg,mean}$, and use it to rescale the image so the scale of intensities is closely maintained. We then subtract $I_{bg,mean}$ to set the background to 0 (i.e., $I_{final} = I_{bg,mean}*(I_0/I_{bg}) - I_{bg,mean}$) Furthermore, one must also choose whether one would like to offset the final result by any nominal values (i.e., $I_{final} = I_0 - I_{bg} + offset$ or $I_{final}$

= $I_{bg,mean}*(I_0/I_{bg})$ - $I_{bg,mean}$ + *offset*). One advantage of doing so is that one can avoid truncating pixel data. If offset is set to 0, then half of the background pixels will be negative, which is problematic for most non-float image formats. It is possible that some image features of interest are below background. Likewise, potentially the pixel-wise noise of the background is of interest and preserving that noise is to advantage. For example, preserving the noise produces a more natural looking final image and eliminates discontinuities or artifacts that may impact subsequent image processing algorithms like derivative calculations. In our case, we chose to use $I_{bg,mean}*(I_0/I_{bg}) - I_{bg,mean} + offset$ as the correction method for immunofluorescence images, setting the offset value to 500 for all datasets, which is ~10 times the pixel-wise standard deviation of the background. This helps to flatten background while retaining the natural noise characteristics of the image, producing a more natural final result without truncation artifacts. For phase images we chose to use $I_{final}=I_0-I_{bg}+offset$ where offset was 0, given values below 0 should not be valid in our cell imaging application. Lastly, since phase shifts are typically expressed in radians and efficient storage of the images requires integer values, and we are performing relative quantification rather than absolute quantification, we scaled the phase image results by a constant factor to maintain data precision within the maximum scale of the saved image.

The general method for calculating image background in Equation 9 has multiple strengths, including speed and flexibility. It is fast given it uses fast linear filtering to compute the final background. It is flexible because it allows local spatial weighting via the averaging kernel $f_k()$ while also enabling a second, completely independent and user-defined approach to discriminating background through the use of binary or gray-scale weighting masks. In our case, we leveraged the flexibility to tailor the background subtraction approach to the different needs of the fluorescent imaging and quantitative phase imaging (QPI). Unlike nearly all other fluorescent image correction methods, the method used here discriminates background based upon pixel-wise variance instead of image intensity, allowing correction in the presence of objects that may absorb light (e.g., red-blood cells) or fluoresce (e.g., Hoechst labeled cells), avoiding artifacts that absorbing particles can impart on intensity-based subtraction methods. In the case of QPI, it allowed us to combine two methods of identifying background pixels to create a more robust background subtraction approach.

*Efficient binary mask generation*

The method of background calculation is closely tied to the process of generating masks for single-cell quantification. In other words, if one can reliably generate a weighting mask for background pixels, one has also likely defined an approach to generating masks for identifying objects of interest (e.g., cells). Since the phase intensity is related to physical quantities of thickness and index of refraction, we were able to set a uniform threshold of 0.25 radians across datasets. Although not utilized here, we found that the standard deviation filtered image was very useful for thresholding fluorescent images when needed. Dividing the original image by the standard deviation image creates an image that represents signal-to-noise ratio. Therefore, for a vast majority of fluorescent imaging applications, one can uniformly set a masking threshold based upon a statistical justification that objects represent signals multiple standard deviations above background (e.g., 5σ) relative to background noise. Thus, in many ways, weighted mean background subtraction method is performing two tasks simultaneously: (i) background subtraction to generate images for quantification, and (ii) binary mask generation to identify quantification on a per-object basis. This dual purpose further enhances the overall computational efficiency of the workflow.

*Fluorescence intensity compensation*

Upon quantifying the individual cells over time, it becomes clear that the limited cell permeability of the Hoechst fluorescent prevents the dye from entering cells instantaneously. Therefore, one observes an asymptotic process of dye entering the cells that starts quickly but takes time to approach steady-state. Therefore, when attempting to apply thresholds of Hoechst fluorescence to determine if a cell has been compromised, either the threshold should be adjusted over time, or “compensated” version of the Hoechst intensity data could be calculated to account for this effect. We chose to perform the latter. Furthermore, we chose to compensate data by only applying an offset to intensity values instead of a scale and offset as the latter proved too difficult to solve robustly for all timepoints across all experiments and experimental conditions.

To find the necessary offset value for each timepoint, we first needed to estimate how much the intensity shifts over time. To do that, we utilized density histograms of the fluorescence data. Timepoints were sequentially grouped into periods of 4 timepoints each (i.e., period1={1, 2, 3, 4}, period2={5, 6, 7, 8}, …) for generating density histograms. Timepoints are acquired every 30 minutes, meaning each period represents 2 hours. Creating histograms from 4 timepoints at a time, provides more datapoints for robust/smooth histogram generation while still retaining time-resolution. Density histograms were made from each period using bin widths defined using the *bw.nrd0()* function in R.

We start the process of aligning these histograms by first defining a “reference” distribution. In this process we start with the distribution associated with the last period of data. The distribution for the previous period then becomes the “test” distribution that will be aligned to the reference distribution. Once the test distribution is aligned, we make the current test distribution the reference distribution, and assign the distribution from the previous period as the test distribution and repeat the alignment process.

Each histogram is aligned by offsetting the $x$ axis (i.e., Hoechst fluorescent intensity) and minimizing an error function. The calculated error was the linear addition of two weighted penalties. The penalty for horizontally shifting the test distribution relative to the reference distribution was weighted by a factor of $\alpha = 0.1$ while the penalty associated with differences in the $y$-values of the test distribution relative to the reference distribution was weighted by a factor of $\beta = 1$. Doing so allows reasonable freedom to shift the test distribution to better align the peaks of the two distributions while preferring as small a horizontal shift as possible. Furthermore, the penalty function utilized a bias parameter, $b$. When $b = 0$, the values of the distribution were uniformly weighted across the $x$-axis. When $b \neq 0$, the cumulative density function of the reference $y$-values ($cdf(y_{ref})$) was used to weight the error, to either favor alignment of distribution peaks further left (b < 0) or further right ($b > 0$) in the distribution. In the case of the Hoechst fluorescent data, the lower distribution represents the live-fraction of cells. Therefore, we used the left-biased formulation by setting $b = -1$. Note that because the test distribution is shifted relative to the reference distribution, differences between the test and reference distributions can only be calculated where the x-values overlap. For this same reason, the cdf is only calculated on this overlapping interval. Also note that the error calculations had to utilize normalized representations of the horizontal and y-value penalties to avoid penalties increasing or decreasing significantly based solely on the number of overlapping datapoints. Thus, the horizontal shift, *d*, was normalized by the total range of the test *x*-values, *L*. Normalization of y-error penalty required two steps. First, the differences in *y*-values between the test ($y_{test}$) and reference ($y_{ref}$) distribution were normalized by the maximum test *y*-value, $y_{max}$. Second, the *average* sum-square-error of the normalized differences was calculated by dividing the sum square error by the number of overlapping datapoints, *n*. The final error, *E*, or penalty associated with a particular distribution shift, *d*, was then given by the following equation.

$$E = \alpha \left(\frac{d}{L}\right)^2 + \beta \frac{\sum \left(\gamma \frac{y_{test} - y_{ref}}{y_{max}}\right)^2}{n} \tag{10}$$

$$\gamma = \left\{ \begin{array}{ll} 1 & \text{for } b = 0 \\ cdf(y_{ref}) & \text{for } b > 0 \\ 1 - cdf(y_{ref}) & \text{for } b < 0 \end{array} \right\} \tag{11}$$

The *gridSearch* function of the *NMOF* package for R was used to search shifts that range 0% and 50% of the total range of x-values observed over the entire timelapse with excess resolution to reliably determine a global minimum in the cost function within the realistic range of shifts.

*Semi-automated threshold determination*

To make data analysis more objective, we also sought to automate determination of the two key thresholds used to quantify cell viability, the Hoechst nuclear staining threshold and the phase image threshold. To do this, the phase and compensated fluorescence data were standardized. For phase data, this was done by dividing the phase intensity by the peak of the density histogram (i.e., an estimate of the mode), subtracting 1, and then multiplying by 10 (for convenience of scale). For fluorescence data, standardization was done by subtracting the median and dividing by the median absolute deviation (MAD) estimate of the standard deviation.

With the data standardized to a predictable scale, a threshold discriminating the live-cell and dead-cell population was predicted two different ways. First, a two population Gaussian mixture model was fit to the distribution using the *emcluster* function of the *EMCluster* package for R. The threshold is defined as the boundary between the two predicted clusters. The second method utilizes a heuristic method built upon the observation that the live cell population is a largely symmetric and Gaussian-like distribution that slightly overlapped the dead-cell population on one side. The standard deviation of the live-cell population was estimated by cutting the distribution at the live-cell population peak, keeping the non-tailed half, then reflecting that non-tailed portion to estimate a full symmetric distribution, and then calculating the MAD estimate of the standard deviation. From that MAD, we could choose a threshold that was a certain number of standard deviations from the live-cell peak. For fluorescence, that meant choosing the live-cell population mode plus 1.64*MAD and minus 1.64*MAD for phase signal given that increases in fluorescence indicate death while decreases in phase indicated death. The semi-manual aspect of the process was that the user would look at the plot of the distribution and the lines that indicated the two estimates of a threshold and decide which of the two estimates provided better captured 95% of the live-cell population. The *emcluster* algorithm generally worked better for the fluorescence data while the MAD-based calculation worked better for phase.

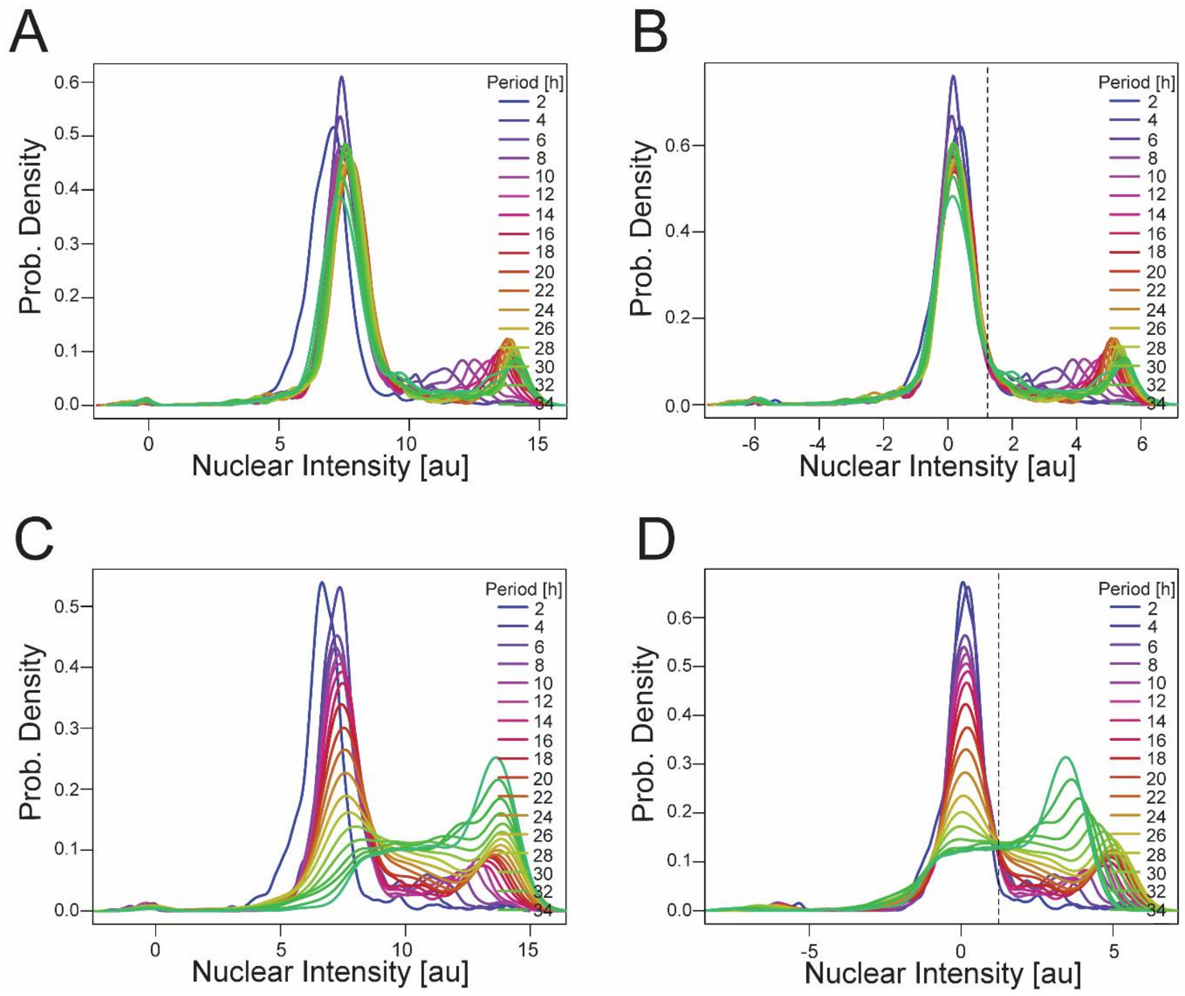


**Supp. Figure 2** – Example of results of fluorescence compensation, standardization, and automated thresholding (*emcluster* algorithm). (A) Density histograms of nuclear staining intensity for untreated cells *before* fluorescence compensation for slow uptake of Hoechst 33258 dye. (B) Density histogram of standardized nuclear staining intensity for untreated cells *after* fluorescence compensation. One can see how the left-hand cluster of cells representing the live cells increases fluorescence intensity over time. After compensation, this drift is removed. This allows a threshold to be drawn at the right edge of the live-cell population for more sensitively and accurately detecting when cells become compromised. (C-D) Density histograms before and after compensation when treated with 5 nM Bort. Again, the compensation removes drift in the live-cell cluster for more accurate application of a threshold to determine cell viability as show in (D).

*JEX Workflow*

Included in the supplemental information is a file ('JEX Workflow Template.txt') that contains all the functions and parameters of the JEX image analysis workflow used for this study. The file can be loaded using the 'Load' button on the 'Process' tab of the JEX software. The functions in the workflow are listed below in order with brief explanation as to their use/function.

1) 'Import Virtual Image Updates' - Create a link to the raw data without having to copy the raw data into the database.
2) 'TIE Phase Calculator' - Calculate the background corrected 'Phase' image object.
3) 'Weighted Mean Filtering' - Calculate the background corrected 'Fluor' image object.
4) 'Merge Objects (virtual)' - Merge the 'Phase' and 'Fluor' image object into a single object without having to copy their image data, making them equivalent to a

multichannel image set with one channel being phase and other nuclear fluorescence.

5) 'Find Maxima Segmentation' - Find the maxima in the phase channel to identify the locations of individual cells. Also generate a mask that defines the “valleys” in-between the cell maxima. The lines defined by these valleys are used to split cell regions of interest in step 9.
6) 'Track Points (LAP)' - Assigns identification numbers (IDs) to each cell that persist over time so that the cell labeled #1 in the image for timepoint #1 is labeled as cell #1 in the image at timepoint 2 (if it is still present). For detailed information on the LAP tracking algorithm used to perform this, please refer to Tinevez et al and Chenouard et al.[4,5] In addition, the function also calculated the median change in position of the cells between each timepoint for the next image registration step.
7) 'Register a Multi-color Image Set (Roi)' - Calculating the cropping rectangles that would be used to create new images that are registered to one another to remove potential inconsistency in the microscope stage. However, just the rectangles are saved instead of new registered images. These rectangles are used in during quantification as the virtual bounds of the image. This avoids unnecessarily duplication of the large image dataset.
8) 'Adjust Image Intensities (Multi-Channel)' - Threshold the phase image to define the cell region of interest for quantifying nuclear staining intensity and phase intensity.

9) 'Prepare Masks for Feature Extraction' - Overlay the cell masks with the valleys mask from step 5 to make sure that each cell mask is associated with 1 and only 1 ID number.
10) 'Feature Extraction' - Quantify the nuclear and phase intensities in the cell mask region within the background subtracted versions of the phase and nuclear fluorescence images.

*Software Code*

Below are links to the repositories and branches used by JEX for this study. The projects are maintained as “Maven” projects and imported as such into the free Eclipse IDE for java developers. Updates and notes on installation are included in the README.md file displayed on the main page of the repository.

JEX - https://github.com/jaywarrick/JEX

Imagej-ops - https://github.com/jaywarrick/imagej-ops

ImgLib2-roi - https://github.com/jaywarrick/imglib2-roi

ImgLib2 - https://github.com/jaywarrick/imglib2

Below are the links to the code used as part of R data analysis.

Functions used to manipulate and plot data.table objects as well as to more easily read data in from JEX repositories - https://github.com/jaywarrick/R-General/blob/master/.Rprofile

Functions used to perform specific single-cell analysis tasks - https://github.com/jaywarrick/R-Cytoprofiling/blob/master/PreProcessingHelpers.R

Below is a link to another repository where key functions built into JEX are provided as ImageJ plugins.

ImageJ-ResearchPlugins - https://github.com/jaywarrick/ImageJ-ResearchPlugins

### Supplemental References


1. Markovina, S. *et al.* Bone marrow stromal cells from multiple myeloma patients uniquely induce bortezomib resistant NF-κB activity in myeloma cells. *Molecular Cancer* **9**, 176 (2010).
2. Zuo, C., Chen, Q., Li, H., Qu, W. & Asundi, A. Boundary-artifact-free phase retrieval with the transport of intensity equation II: applications to microlens characterization. *Opt Express* **22**, 18310–18324 (2014).

**Figure Legends**

**Supp. Figure 1** – Plot of weighting function as a function of *sd(m,n)* for $\delta=10^{-7}$, $h_{mode}=0.3$, $h_{min}=0.1$, and different scaling factor values, *p*. The weighting function is a symmetric around the value of $h_{mode}$, with a maximum value of 1 at $sd(m,n)=h_{mode}$. The function reduces to 0.5 at $sd(m,n)=h_{mode}\pm(h_{mode}-h_{min})$ and rapidly approaches 0 as $sd(m,n)\rightarrow\pm\infty$. Different values of *p* change how quickly weights drop off above and below $h_{mode}$ thereby controlling how sharply weighted background pixels are in the weighted mean calculation. Also notice that the closer the values of $h_{mode}$ *and* $h_{min}$ are to one another, the tighter the weighting scheme is focused around $h_{mode}$. Effective values of p are generally between 0.3 and 5 with a value of 2 being used for our datasets.

**Supp. Figure 2** – Example of results of fluorescence compensation, standardization, and automated thresholding (*emcluster* algorithm). (A) Density histograms of nuclear staining intensity for untreated cells *before* fluorescence compensation for slow uptake of Hoechst 33258 dye. (B) Density histogram of standardized nuclear staining intensity for untreated cells *after* fluorescence compensation. One can see how the left-hand cluster of cells representing the live cells increases fluorescence intensity over time. After compensation, this drift is removed. This allows a threshold to be drawn at the right edge of the live-cell population for more sensitively and accurately detecting when cells become compromised. (C-D) Density histograms before and after compensation when treated with 5 nM Bort. Again, the compensation removes drift in the live-cell cluster for more accurate application of a threshold to determine cell viability as show in (D).

**Supp. Figure 1**

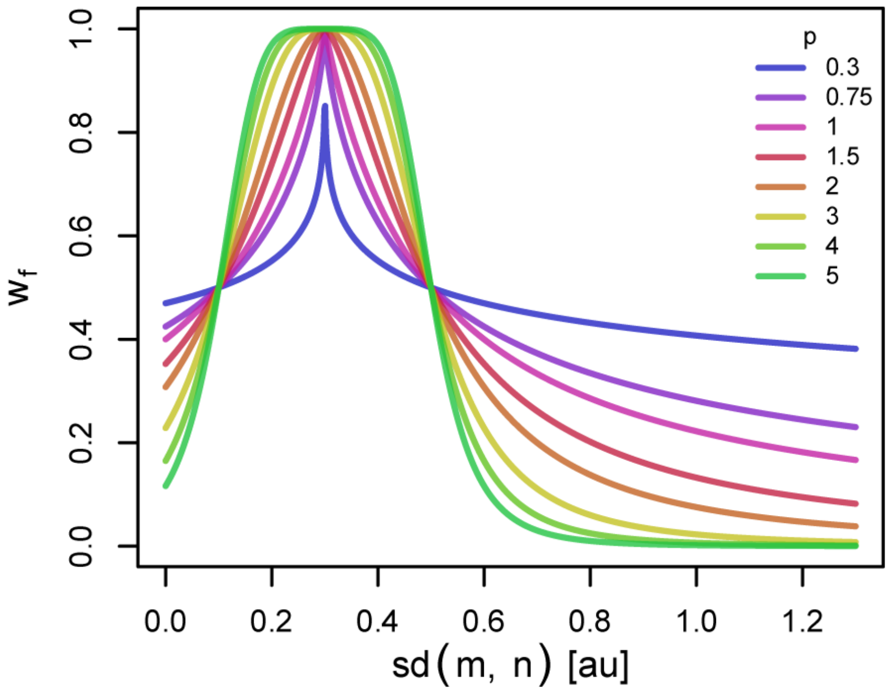

**Supp. Figure 2**

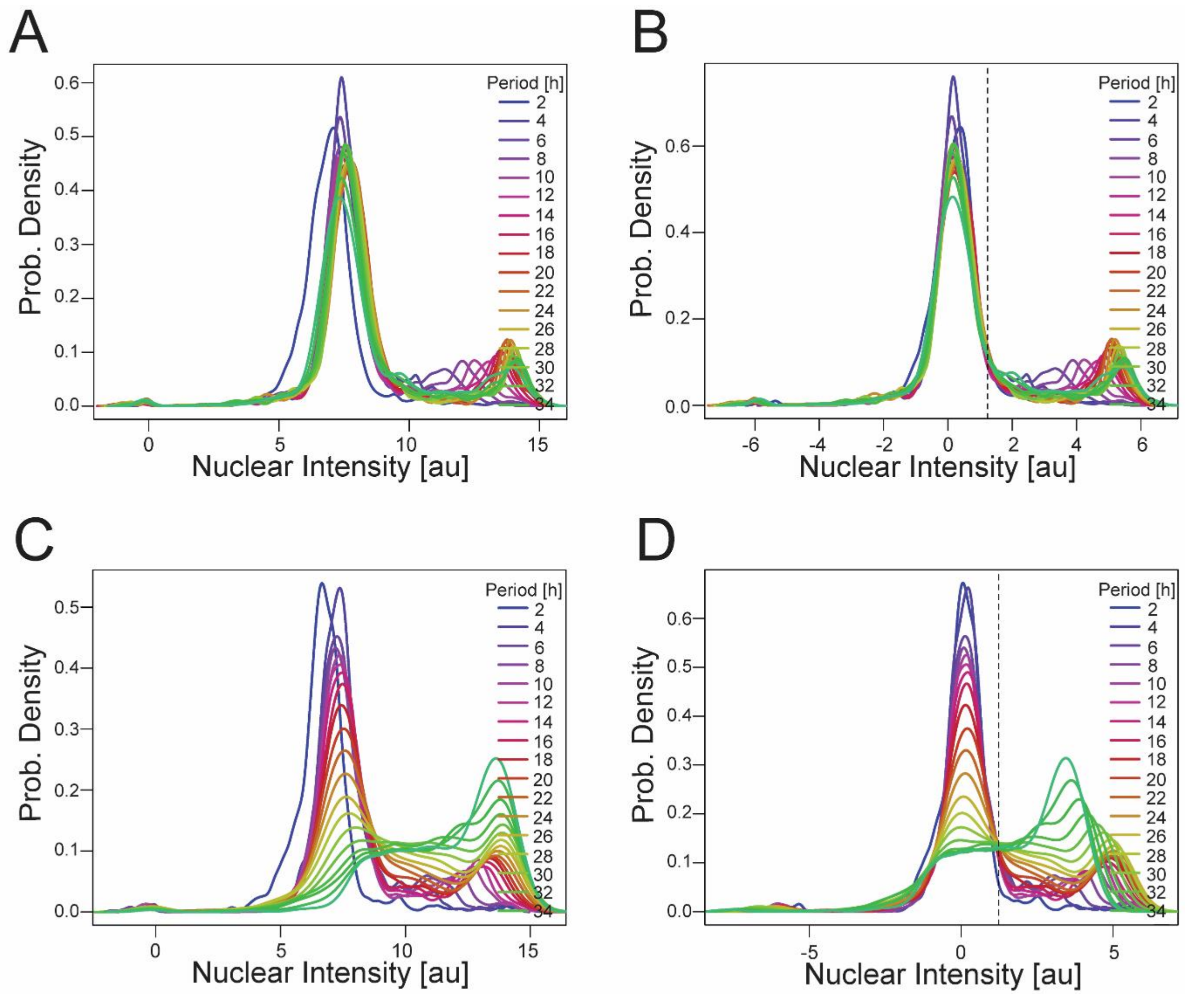
